# Protein fold prediction using simulated DEER distance distributions and decay traces

**DOI:** 10.1101/581181

**Authors:** Diego del Alamo, Maxx Tessmer, Richard Stein, Jimmy B. Feix, Hassane S. Mchaourab, Jens Meiler

## Abstract

Despite advances in sampling and scoring strategies, Monte Carlo modeling methods still struggle to accurately predict *de novo* the structures of large proteins, membrane proteins, or proteins of complex topologies. Previous approaches have addressed these shortcomings by leveraging sparse distance data gathered using site-directed spin labeling and electron paramagnetic resonance spectroscopy (SDSL-EPR) to improve protein structure prediction and refinement outcomes. However, existing computational implementations must choose between coarse-grained models of the spin label that lower the resolution and explicit models that lead to resource-intense simulations. Existing methods are further limited by their reliance on distance distributions, which are calculated from a primary refocused echo decay signal and may contain artifacts introduced during this processing step. Here, we addressed these challenges by developing RosettaDEER, a scoring method within the Rosetta software suite capable of simulating distance distributions and echo decay traces between spin labels fast enough to fold proteins *de novo*. We demonstrate that the accuracy of resulting distance distributions match or exceed those generated by more computationally intensive methods. Moreover, decay traces generated from these distributions recapitulate intermolecular background coupling parameters, allowing RosettaDEER to discriminate between poorly-folded and native-like models even when the time window of EPR data collection is truncated, rendering them unsuitable for accurate transformation into distance distributions. Finally, we demonstrate that one decay trace per nine residues is sufficient to predict the folds of Bax and the C-terminus of ExoU, two soluble proteins with surface-exposed amphipathic structural features that prevent the Rosetta energy function from correctly identifying native-like models in the absence of experimental data. These benchmarking results confirm that RosettaDEER can effectively leverage sparse experimental data for a wide array of modeling applications built into the Rosetta software suite.

## Introduction

Structural biology increasingly relies on integrated methods to model the structure and dynamics of proteins and protein assemblies^1,2^. Multiple complementary experimental methodologies, when integrated with computation, can describe the structure and dynamics of proteins that elude structure determination from a single technique, such as integral membrane proteins, conformationally flexible proteins, and those that fall outside the size limitations of solution-state nuclear magnetic resonance and cryo-electron microscopy. Computational approaches are used to integrate experimental data from multiple approaches and build physically-realistic models also in regions with sparse experimental data. One promising experimental approach to feed into integrated structural biology combines site-directed spin labeling and electron paramagnetic resonance spectroscopy (SDSL-EPR). Previous studies have employed SDSL-EPR and computation in tandem to predict protein structures *de novo* ^3–10^, model conformational changes^11–14^, and dock rigid-bodies^15–17^.

Existing modeling methods largely focus on data gathered using four-pulse double electron-electron resonance spectroscopy^18^ (DEER, also called PELDOR), which can report on distances of up to 60 to 80 Å between stable unpaired electrons conjugated to the protein backbone by SDSL^19,20^. However, incorporation of these distances as interatomic restraints is confounded by the conformational freedom of the most commonly used probe, the methanethiosulfonate spin label (MTSSL). The central challenge is to convert interspin distance information of a mutant protein into backbone restraints for the wild-type^21–23^. The need to incorporate two spin labels into the protein sequence per restraint results in sparse structural coverage of the experimental data that can introduce ambiguities into computational modeling^6^. Only a few experimental restraints are generally available to describe the protein’s conformational details.

These sparse datasets have nonetheless been leveraged for protein structure prediction and refinement by a range of computational modeling approaches that represent the spin labels either implicitly or explicitly. Implicit models such as the motion-on-a-cone (CONE)^3^ model use knowledge-based potentials to translate inter-spin distance values into backbone restraints, typically between C_β_ atoms. Introducing these restraints led to measurable improvements in de *novo* structure prediction benchmarks by programs employing Monte Carlo sampling strategies^3,5–7,23,24^ gradient minimization^4,10^, and molecular dynamics^25^. However, because these potentials account for neither the residues’ surrounding environment nor their relative orientations, they tend to be relatively imprecise^26^. Explicit methods, by contrast, model spin labels as either individual side chains^14,27–30^, ensembles of side chains^15,31–33^, or ensembles of dummy atoms^34^. The added detail improves accuracy of modeling but makes implementations too computationally intensive for *de novo* protein structure prediction and limits the utility of these methods to validating experimental distance distributions^35^ and modeling small-scale conformational changes^11,12,14^.

Despite their diversity, these methods largely share a common limitation in their reliance on distance distributions, rather than the primary spectroscopic readout. Other computational methodologies directly incorporate primary experimental data, such as two-dimensional NMR spectra^36^ and cryo-EM electron density maps^37^ to fold and refine proteins. The feasibility of using DEER dipolar coupling decay traces as modeling restraints, by contrast, has only recently been explored^14^. Whereas processing spectroscopic decay traces into distance distributions risks the introduction of artifacts, simulating a decay trace from a distance distribution is well-described and mathematically straightforward^20^.

Here we introduce RosettaDEER, a method in the macromolecular modeling suite Rosetta capable of rapidly simulating distance distributions and DEER decay traces between spin labels as well as evaluating a model’s agreement with experimental data. RosettaDEER’s increased computational efficiency enables prediction of protein structures *de novo* with greater accuracy than the default energy function or the previously reported CONE model^3^. Owing to Rosetta’s Monte Carlo sampling strategy^38^, the experimental data can be used directly without analysis or background-correction. Thus, the quality of the primary spectroscopic data can be significantly poorer than what would ordinarily be required for rigorous transformation into distance distributions. This method reinforces the utility of DEER in conjunction with computational modeling to accurately model proteins structures.

## Results

### Modeling nitroxide spin labels using RosettaDEER

A strategy to model proteins using DEER data must reliably simulate distance distributions between spin-labeled residues. To quantify the computational cost and efficiency of this task, we considered a panel of five proteins where both atomic-detail structures and experimental DEER data were available (Table S1). Distance distributions were simulated with each method between residue pairs that have been previously studied experimentally, and the resulting error was quantified as the difference between the average values of the simulated and experimental distance distributions (Figure 1A). In addition, we measured the time required by each program for in *silico* spin-labeling of a single residue (Figure 1B). Consistent with previous results^8,26,32^, the average values of experimental distance distributions gathered in monomeric proteins, but not the homodimer CDB3, agree more closely with those of simulated distributions than their corresponding C_β_-C_β_ distances, from which restraints such as the CONE model are derived^3^ (Table S2). By contrast, none of the methods examined here reliably reproduced the width of the experimental distributions. This is likely attributable to oversampling of available conformational space of the spin label, which results from the exclusive use of van der Waals repulsive energies to limit side chain configurations and, by extension, electron positions for distance measurements. Finally, the data revealed how simulation times both varied substantially between these methods and failed to correlate with accuracy.

**Figure 1.**
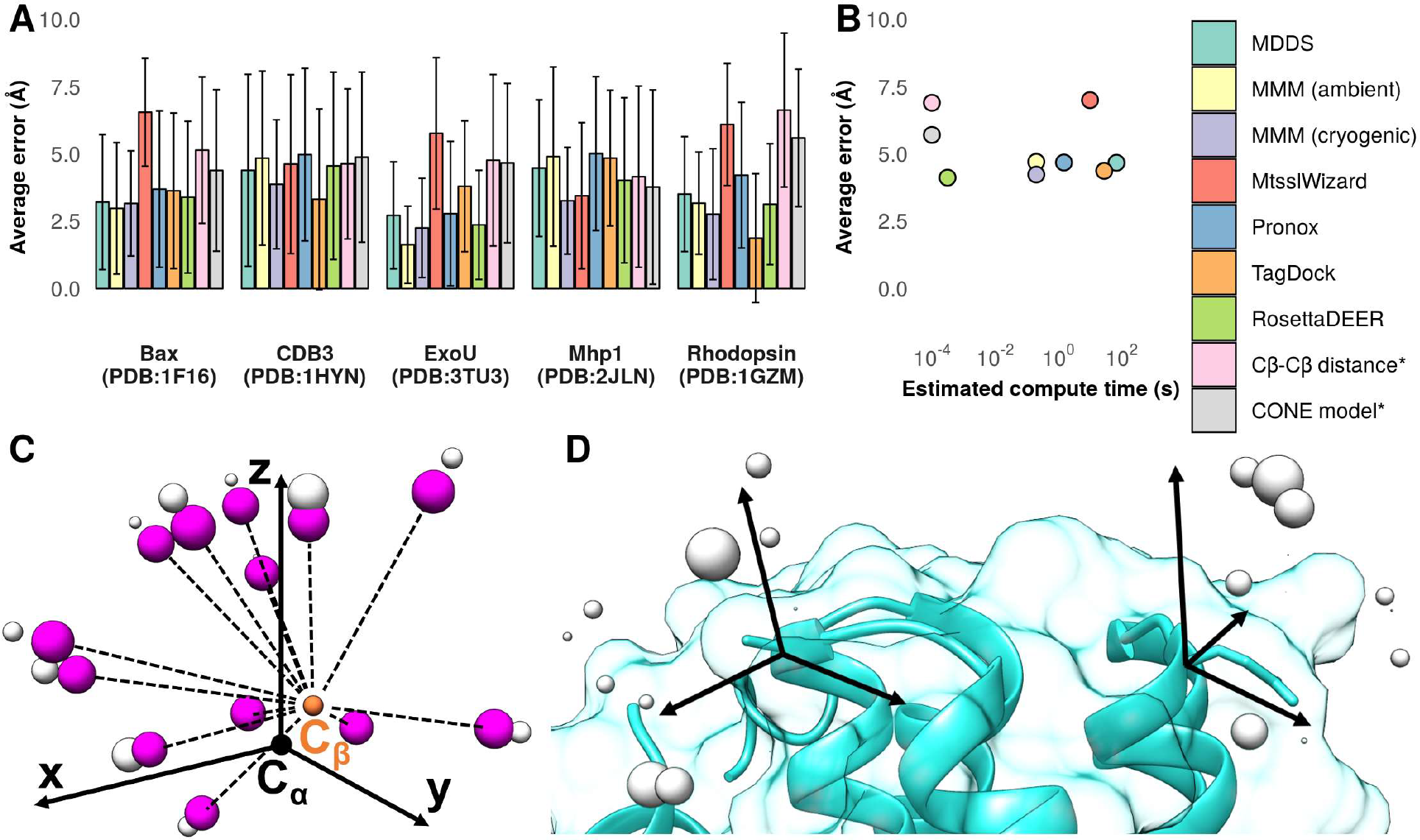
Simulations of distance distributions between nitroxide probes using RosettaDEER. **A)** Difference between average values of simulated and experimental distance distributions. Error bars reflect the standard deviation. B) Estimated average time required for in silico spin labeling of one residue (*the lower limit of quantitation exceeded the C_β_-C_β_ distance compute time). **C)** Coarse-grained rotameric ensemble representation of MTSSL. Centers of mass, shown in purple, are used for clash evaluation, while electron coordinates, shown in grey, serve as measurement coordinates. D) Distance distributions between residues are simulated by superimposed coordinates, evaluating clashes, and measuring all resulting pairwise distances

Because these results suggested that computational complexity was not a determining factor in the accuracy of simulations of distance distributions between spin labels, RosettaDEER’s design prioritized computational speed over accuracy (see Methods). Rather than measure distances from full-atom rotamers or mobile dummy atoms, RosettaDEER instead uses a probability density function to capture the likely positions that would be explored by MTSSL conformations. High-occupancy electron positions are mapped on the protein structure (Figure 1C, 1D). For each electron position, a clash evaluation was performed between a centroid atom representing the nitroxide ring’s center of mass and the protein backbone. Placing this coordinate at an idealized location, consistent with spin-labeled protein structures in the Protein Databank (Figure S2, Table S3), reduced the number of atoms for clash evaluation to one per rotamer, thus maximizing computational efficiency. Figures 1A and 1B demonstrate that, compared to other methods, RosettaDEER’s simplified representation of the spin label allows the generation of distance distributions three to five orders of magnitude faster than other approaches but with comparable accuracy.

### Comparison of simulated with experimental DEER decay traces

We then explored approaches to simulating DEER decay traces using these distance distributions, a task complicated by the fact that experimental decays have contributions from coupling between unpaired electrons both within and between macromolecules^20^ (Figures 2A-C). Model-free analytical methods such as Tikhonov regularization rely on appropriate time collection windows to isolate, and thus correct for, the intermolecular signal. Our approach instead borrows from Gaussian-based modeling approaches that fit the data directly by treating the intermolecular contribution as an exponential function consisting of a slope (*k*, the background decay) and a y-intercept (*λ*, the modulation depth; see Methods)^39^. We tested this strategy on our benchmark set and found that both parameters agreed with experimentally determined values (Figures 2D, 2E), with r^2^ values of 0.92 and 0.95 for k and *λ*, respectively. Perhaps unsurprisingly, the outliers in this respect tended to be the decay traces with the fewest oscillations (Table S4, Figure S3). This functionality enables RosettaDEER to directly evaluate structural models while avoiding the shortcomings associated with transformation to distance distribution.

**Figure 2.**
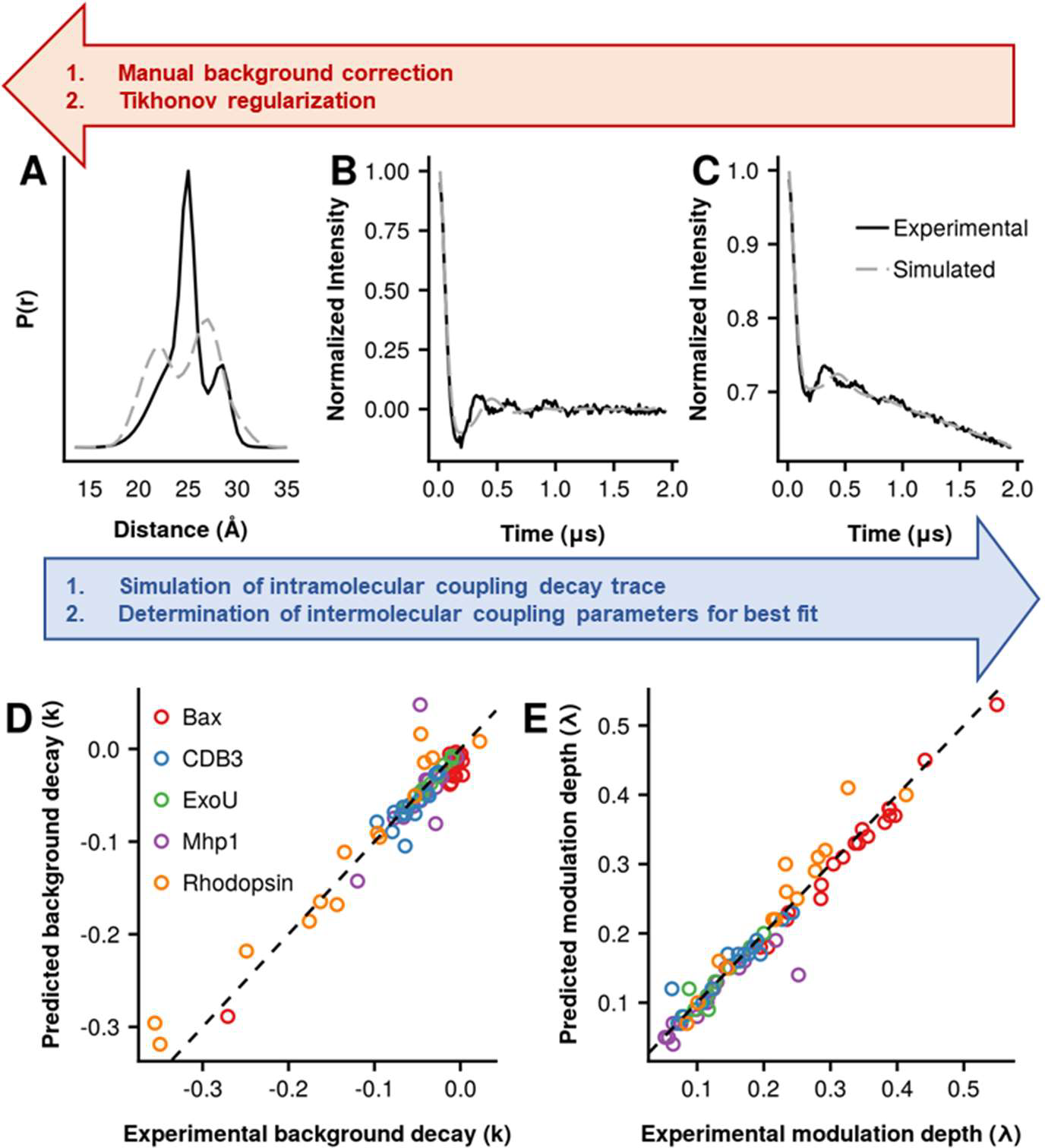
RosettaDEER simulations of distance distributions and decay traces. The forward approach taken by RosettaDEER contrasts with the ill-posed inverse problem that must be addressed by Tikhonov regularization. **A)** Simulated and experimental distance distributions between T4 Lysozyme residues 93 and 123. **B)** RosettaDEER then simulates the resulting intramolecular decay trace and determines the background parameters, *k* and *λ*, which result in the generation of a trace that best fits the experimental data (**C**). **E and F)** Recovery of experimental background coupling decay parameters.

### Enrichment of native-like models using experimental decay traces

The extent to which the added capability of directly simulating decay traces could improve the identification of correct protein structural models was evaluated using misfolded and misdocked structural models. We generated one to two thousand incorrectly folded models for each of the proteins mentioned above. In addition, we generated one thousand misdocked models of the homodimer CDB3 (Figure 3). The root mean square deviation over secondary structures normalized to a 100 residue protein (RMSD100SSE^40^) ranged from near-native models (0.5 Å C_α_ RMSD100SSE) to incorrectly folded (>15 Å C_α_ RMSD100SSE). To assess enrichment of high-quality models, we calculated the logarithm of the number of models in the top ten percentile by agreement with DEER data that also fell in the top ten percentile by RMSD100SSE as previously described^7^ (see Methods). This resulted in a metric ranging from −1 (no high-quality models in the top 10%) to 1 (only high-quality models in top 10%), with a value of 0 indicating no enrichment. This scoring scheme was repeated for the native Rosetta energy function^41^ and the CONE model (Figure S4). To examine the synergistic effect of using both the energy function and either RosettaDEER or the CONE model, the combined Z-scores were compared for each model and plotted to highlight the level of enrichment (Figure S4).

**Figure 3.**
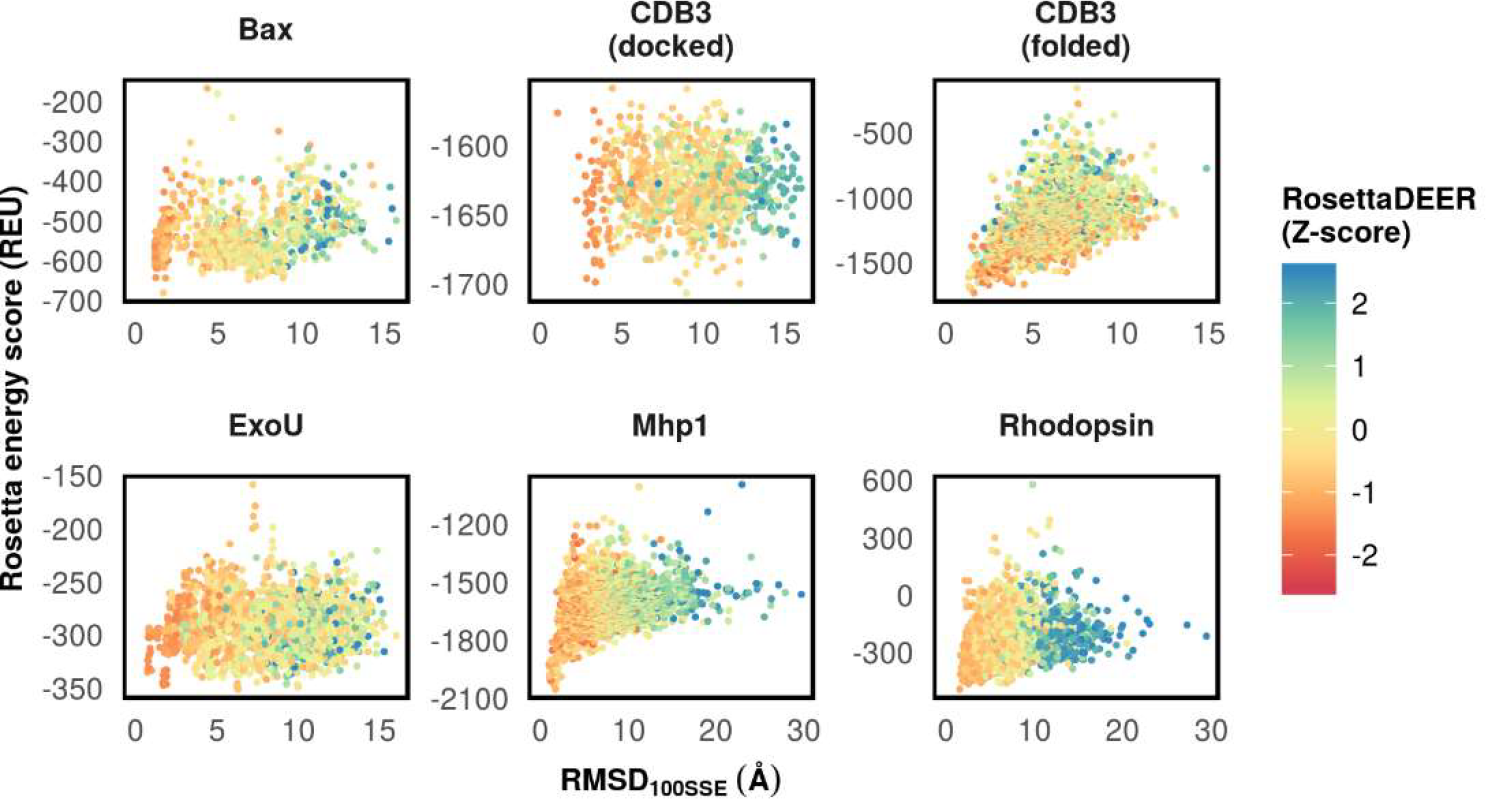
Evaluation of models using DEER decay traces. Models with C_α_ RMSD100SSE ranging from 0.5 Å to 20.0-30.0 Å were scored using both the Rosetta energy function and RosettaDEER.

For the monomeric proteins considered here, decay traces simulated from native-like models more closely corresponded to experimental data than those from incorrectly folded models. In all cases examined, RosettaDEER identified correctly folded models as well as correctly-docked CDB3 models more effectively than the CONE model and the Rosetta energy function on its own. In contrast, misfolded models of CDB3 were not identifiable using either RosettaDEER or the CONE model with the experimental data available. In fact, the use of the experimental data alongside the Rosetta energy function impeded the latter’s identification of native-like models. This may reflect the fact that the experimental data gathered in CDB3 is limited to interdimeric distance restraints, which reflect the relative position of a residue from the center of symmetry, rather than structural features within the protein fold.

Because a robust simulation of the intermolecular background is dependent on the time window of experimental DEER trace collection, we examined the influence of the decay duration on the enrichment of correctly-folded protein structural models (Figure S4). Model-free analytical approaches require approximately 0.8 oscillations and 1.6 oscillations to accurately identify the average and standard deviation of a distance distribution, respectively^20^. Therefore, we artificially truncated the experimental data and measured enrichment as a function of oscillations (see Methods, Figure S4). Strikingly, with RosettaDEER highly truncated decay traces (<0.5 oscillations) could still identify correctly folded models of Bax, ExoU, Rhodopsin, and Mhp1, albeit to a reduced degree. In these cases, as well as for misdocked CDB3 models, enrichment increases with duration up to approximately one oscillation and largely plateau thereafter (Figure S4).

### *De novo* folding of Bax and ExoU

To further illustrate RosettaDEER’s capability to identify native-like models, we folded Bax and ExoU *de novo.*. Few of the ten thousand structural models generated using Rosetta without experimental restraints resembled the native fold. Moreover, native-like models did not correspond to their respective global energy minima as judged by the Rosetta energy function. (Figure 4A). Therefore, we supplemented this *de novo* fold prediction protocol with one experimental restraint per nine residues using either the CONE model or RosettaDEER (Figure S5). Whereas using the CONE model to transform the distances into restraints led to a measurable improvement in the number of correctly folded structural models (<7.5 Å C_α_ RMSD100SSE) of Bax, no such improvement was observed when folding ExoU. By contrast, using these restraints with RosettaDEER substantially increased the number of correctly folded models of both proteins (Figure 4A).

**Figure 4.**
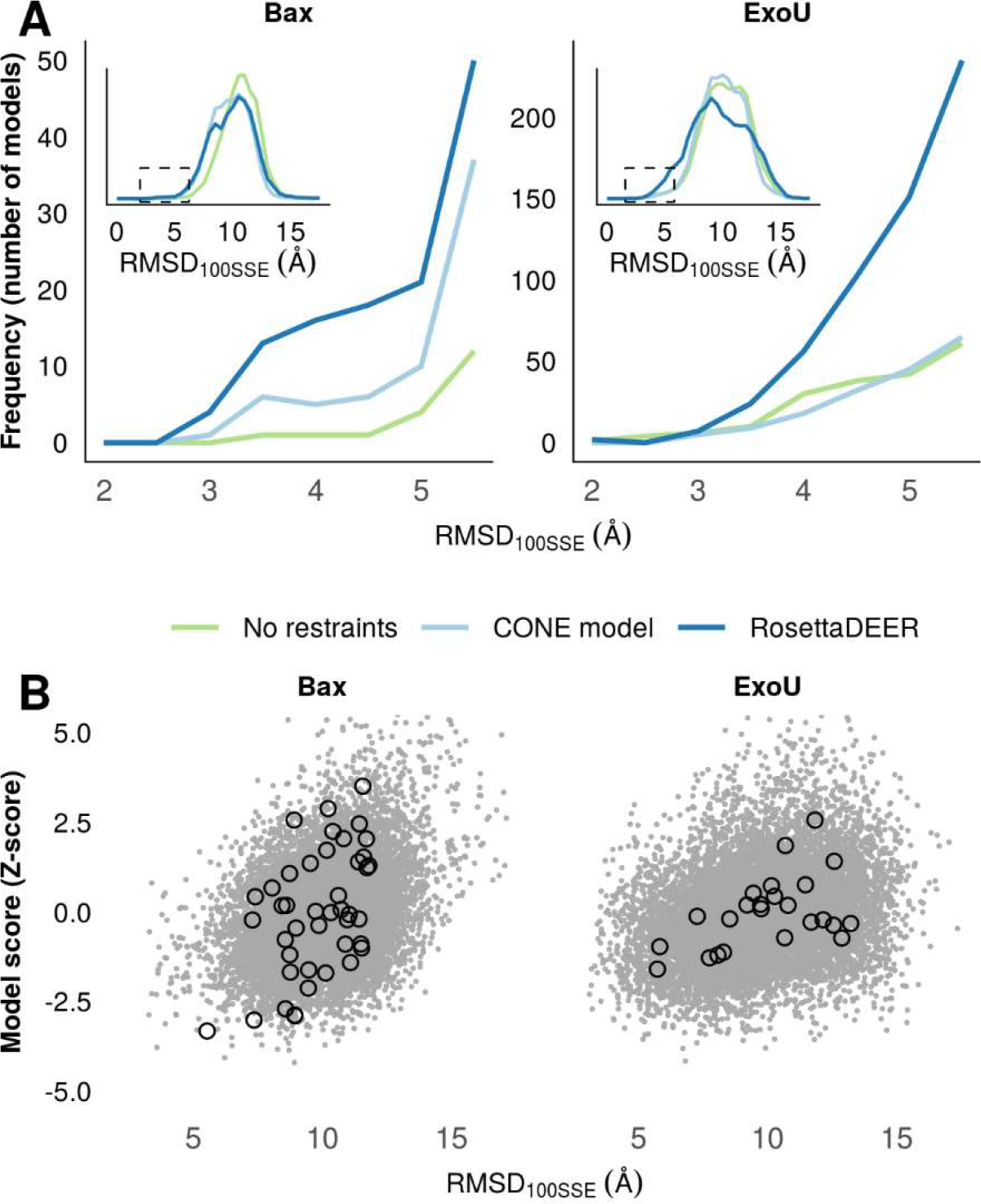
De novo folding of Bax and ExoU using DEER decay data. **A)** De novo protein folding of native-like models using DEER decay restraints with RosettaDEER, C_β_-C_β_ distance restraints with the CONE model, or no restraints. Inset: Spread of all models generated using these three methods. **B)** Accuracy of de novo folded models (gray dots) and clusters (black circles) as a function of combined DEER and Rosetta Z-score.

Although agreement between models and experimental structures loosely correlated with both RosettaDEER score and Rosetta score for both proteins, an abundance of incorrectly-folded models obscured this trend (Figure 4B; RosettaDEER and Rosetta scores were jointly considered by adding the Z-scores of each). As a result, the Rosetta energy function, RosettaDEER, and the CONE model all failed to isolate and identify native-like models for either Bax of ExoU from score values alone. The ten best-scoring models by these metrics were generally incorrectly folded (5-10 Å C_α_ RMSD100SSE) and buried amphipathic features found on the surface of the native model.

This shortcoming was addressed by clustering all models with a radius of 7.5 Å C_α_ RMSD100 using Durandal^42^ and evaluating the size of each cluster and their average agreement with both the experimental DEER data and the Rosetta energy function. Correctly-folded protein models have previously been identified near the center of large clusters^43^, suggesting an additional venue to identifying correctly folded models. To test this hypothesis, we computed the Z-scores of both the average Rosetta energy score and RosettaDEER score for models within each cluster and ranked the clusters by their combined Z-scores. The cluster with the lowest average combined Z-score for both proteins also had the lowest average RMSD100SSE (Figure 4B). In both cases, the cluster center was substantial closer than its average (data not shown). Finally, selecting for the top five clusters with the lowest combined Z-scores eliminated 85.3% of Bax models and 61.3% of ExoU models while maintaining the majority of correctly folded models.

Each cluster at this stage represented a broad population of models that satisfied the DEER restraints and were considered energetically favorable by the Rosetta energy function. We hypothesized that refining models in the absence of restraints would reveal the native fold by allowing false positives to be structurally optimized away from conformations consistent with the data. For this purpose, models from the top five clusters were recombined using a recently published refinement protocol^44^. During each of five iterations, a subset of twenty input models were hybridized in the absence of DEER restraints into 240 models using RosettaCM^45^. The models generated this way were then clustered and analyzed as previously described. Consistent with our prediction, the most native-like cluster retained its agreement with the DEER data, whereas the others refined toward conformations inconsistent with the experimental data. After further Cartesian minimization^46^, the best-scoring model from these clusters had near-native folds (<3.5 Å C_α_ RMSD100SSE; Figure 5).

**Figure 5.**
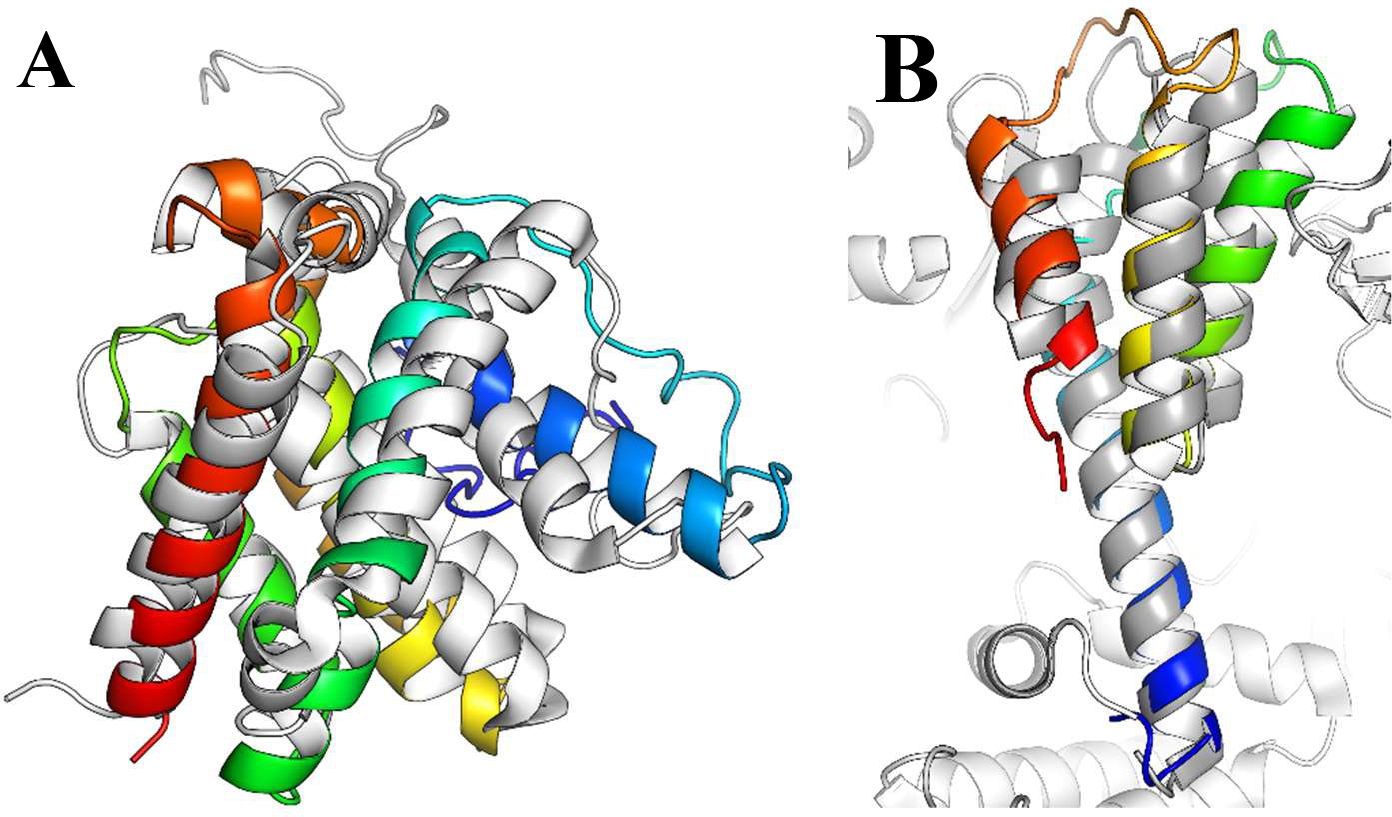
Predicted models of A) Bax and B) ExoU at 3.2 and 2.1 Å C_α_ RMSD100SSE, respectively. Models were obtained from ten thousand *de novo* folded models, the best-scoring of which were refined into 1200 additional models. Native models shown in white.

## Discussion

RosettaDEER predicts and refines protein structures by integrating DEER spectroscopy data and Rosetta computational modeling. The novel aspects of this method are a simplified representation of MTSSL and a strategy to rapidly simulate DEER decay traces for comparison to uncorrected experimental traces. The robustness of the method was demonstrated by benchmarking every step on five sparse datasets. Despite the simplified spin label representation, the distance distributions simulated by RosettaDEER are comparable to those generated using more computationally complex rotamer library approaches. Moreover, despite the imperfect fit to the experimental distance distributions and the decay traces, the combined approach efficiently identifies conformations that simultaneously satisfy the data and the Rosetta energy function.

Our *de novo* folding benchmark with the small soluble proteins ExoU and Bax highlights the success of this strategy. Both proteins possess surface-exposed amphipathic substructures that insert into the membrane. Bax transitions from a soluble monomer into a membrane-bound oligomer using its C-terminal helix^47^, whereas ExoU is hypothesized to move into the membrane using a flexible loop between its two C-terminal helices^48^. Consistent with previous results^8,9^, the Rosetta energy function favored models that packed these substructures in the protein core, leading to incorrectly folded models and lack of correlation of the Rosetta score with model accuracy. As a result, orthogonal experimental data that define the structure are critical to de novo folding. Our benchmark suggests that even a small number of experimental DEER measurements is sufficient to achieve fold-level accuracy. Moreover, our model analysis suggests that even low-quality data can discriminate correctly-folded from incorrectly-folded models more effectively than the C_β_-based CONE model.

The strategy of RosettaDEER to predict the structures of these two proteins (Figure S6) leverages the experimental data by folding and optimizing protein structures with and without restraints, respectively. The first step leads to a substantial reduction in the search space and a concomitant increase in models that satisfy the restraints, although not all of these models are correctly folded. After clustering the models to remove those that correspond to narrow energy minima, the second step, optimization without restraints, allows clusters with incorrectly folded models to adopt alternative folds. This is an effective filtering procedure that restores the experimental data’s ability to identify native-like models. Although the most correct models of Bax and ExoU at this stage were not the best-scoring, they were the most consistent with the experimental data. This protocol therefore minimizes both the generation of incorrectly-folded structures that overfit that data and the conformational search space inherent to the protein folding problem.

Despite its success illustrated here, the current implementation of RosettaDEER’s assumes that a single conformation describes the data. For example, the distance distributions of Mhp1, the most conformationally flexible protein examined in this dataset, were generally more poorly simulated using available methods than those collected in other proteins. Experimental applications of the DEER technique often focus on monitoring ensembles of protein conformations and require computational methods that interpret this data with the capability to generate multiple models and examine their consistency with sparse experimental data. This is the next step of the development of RosettaDEER. Further, a Rosetta *de novo* folding protocol for membrane-associated proteins that includes a model membrane would be desirable for proteins such as Bax and ExoU.

## Methods

### Assembly of diverse experimental datasets

RosettaDEER was implemented in the Rosetta software suite^38^ (Figure S6), trained on distance data gathered in T4 Lysozyme obtained from the laboratory of Hassane S. Mchaourab, and tested and cross-validated using both raw spectroscopic and analyzed distance data gathered in five laboratories. Data for the ExoU C-terminus, Bax, and Mhp1 were obtained from and analyzed by the laboratories of Jimmy Feix, Enrica Bordignon, and Hassane S. Mchaourab, respectively; previously unpublished ExoU double-cysteine mutants were purified, spin labeled, and measured as previously described^9^. Raw data for CDB3 and bovine rhodopsin were obtained from the laboratories of Albert Beth and Wayne Hubbell, respectively, and were analyzed using DEERAnalysis2016^49^. The last 200ns and 500ns were removed from experimental decay traces shorter and longer than 1.5 us, respectively.

### Generation of DEER distance distributions

Distance distributions were obtained from a variety of methods on Bax (PDB: 1F16 model 8), ExoU (PDB: 3TU3), CDB3 (PDB: 1HYN chains R/S), Rhodopsin (PDB: 1GZM chain A), and Mhp1 (PDB: 2JLN) using MMM^31^, MDDS^34^, MtsslWizard^32^, Pronox^33^, and TagDock^16^. MMM2017 was run locally on both cryogenic mode (175 K) and ambient mode (298 K) using default settings. MDDS was run using the CHARMM-GUI web server^50^ on default settings. MtsslWizard was run locally from Pymol 1.7.2.1 using tight fitting unless no rotamers could be placed, in which case loose fitting was used (because Mhp1 residue 324 could not be labeled using loose fitting, distances between it were omitted). Pronox was run from the USC web server using a bias of 0.9 and a van der Waals radius scaling factor of 0.75, the latter of which was reduced to 0.4 if rotamers could not be placed. TagDock was run locally with SCWRL4 and a bump radius of 0.85. Measurements using the CONE model were determined by adding 1.79 Å to the C_β_-C_β_ distance.

### RosettaDEER design

The Rosetta MTSSL rotamer library^29^ served as the basis for the coarse-grained rotameric ensemble used in this study. Each rotamer’s nitroxide bond midpoint was transformed into a common coordinate frame defined by the C_α_ atom at the origin, the backbone nitrogen along the Z-axis, and the backbone carbonyl carbon in the X-Y plane. Centroid atoms with a van der Waals radius of 2.2 Å representing the nitroxide ring center of mass were placed at 87.5% of the distance between each electron and an idealized C_β_ coordinate. Electrons whose respective centroids clashed with backbone atoms and other side chains were eliminated and downweighted, respectively. Inter-electron distances were transformed into Gaussian distributions with bin sizes and standard deviations of 0.5 Å and amplitudes equal to the product of the respective electron weights. All distributions generated this way were added to generate a simulated distance distribution for comparison to experimental values and simulation of DEER decay traces.

Coordinate positions and weights were optimized using a training set of forty-nine experimental distance distributions between 37 residues in T4 Lysozyme^34^. Coordinates were clustered into between two and 53 coordinates using K-means clustering and superimposed over spin-labeled residues. During each of half a million iterations, the weight of an electron was randomly modified and either accepted or rejected using a Monte Carlo Metropolis criterion based on the resulting distributions’ improved agreement with experimental distributions. One thousand repeats were performed for each cluster size. The electron set that led to the greatest agreement with experimental distributions contained 17 coordinates, four of which were downweighted to zero. This set was introduced as the default for RosettaDEER and used for subsequent experiments.

### Simulation of DEER dipolar coupling decay traces

The intramolecular form factor *(Vintra(t))* was generated from each 0.5 Å bin of a distance distribution between 15 Å and 100 Å:

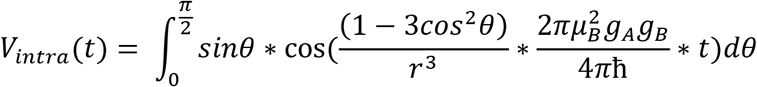

where *t* is the time point of a trace in μs, *r* is the bin distance in nm, *μΒ* is the Bohr magneton, *gX* is the G-factor of electron X, and *θ* is the angle between the interelectron vector and the bulk magnetic field. The intermolecular background consists of the modulation depth (*λ*) and an intermolecular coupling parameter (*k*). These parameters were determined using linear regression by incrementing *λ* with step size 0.01 and linearizing the remainder of the signal:

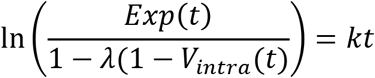

where *Exp(t)* represents the experimental data at time point *t* provided by the user.

### Model generation and evaluation

Diverse sets of models were generated with RosettaCM^45^ using either full-length or truncated native models as inputs. Additional low-quality models were generated using *de novo* protein folding. Bax, ExoU, and CDB3 were scored using the ref2015 energy function^41^, and Rhodopsin and Mhp1 were scored using RosettaMembrane^51^. The transmembrane regions for Rhodopsin and Mhp1 were predicted using OCTOPUS^52^. Oscillation frequencies of decay traces for distributions with an average distance *r*_*avg*_ were calculated as 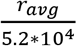 μs^53^. Decay traces with fewer than three oscillations were not used to evaluate enrichment as a function of decay trace duration.

### *De novo* protein structure prediction

Ten thousand models were generated using extended chain AbInitio with either RosettaDEER restraints, CONE model restraints, or no restraints. This protocol relies on insertion of fragments obtained from a July 2011 database; homologous protein structures were excluded from these fragment libraries. The Z-scores of each model’s agreement with DEER data and energy function score was calculated, and all models were clustered to 7.5 Å C_α_ RMSD100 using Durandal^42^. After eliminating sparsely populated clusters (<5% the size of the largest cluster), the top ten models from each of the top five clusters by combined Z-score were iteratively hybridized for five iterations as previously described^44^ without RosettaDEER restraints. The models generated this way were again clustered at 7.5 Å C_α_ RMSD100, and the best-scoring model from the cluster with the best average combined Z-score was treated as the predicted model.

## Author Contributions

DDA, HSM, and JM conceived the study and wrote the manuscript. DDA developed and implemented RosettaDEER and performed computational experiments with input from RAS. MHT and JBF designed and performed DEER experiments on ExoU.

## Acknowledgements

Research was funded by the National Institutes of Health (R01 GM080403, R01 GM073151, R01 GM114234, R01 HL122010, and R01 HL144131). We would like to thank Dr. Christian Altenbach, Dr. Enrica Bordignon, and Dr. Eric Hustedt for providing experimental data used in this study. We thank Dr. Rocco Moretti, Dr. Axel Fischer, and Dr. Andrew Leaver-Fay for helpful discussions on designing and implementing RosettaDEER.

