## Supplemental Data for "Protein fold prediction using simulated DEER distance distributions and decay traces"

**Table S1. Benchmark set for evaluation of RosettaDEER.**

| Protein | Source | Restraints | PDB | Publication |
|---|---|---|---|---|
| **Bax** | Jeschke, ETH Zurich | 21 | 1F16 H | Bleicken (2014) *Mol Cell*[47] |
| **ExoU** | Feix, Med Coll Wisconsin | 11 | 3TU3 | Fischer (2017) *ACS Omega*[9] |
| **CDB3** | Beth, Vanderbilt | 15 | 1HYN R/S | Zhou (2005) *Biochemistry*[54] |
| **Rhodopsin** | Hubbell, UCLA | 14 | 1GZM A | Altenbach (2008) *PNAS*[55] |
| **Mhp1** | Mchaourab, Vanderbilt | 18 | 2JLN | Kazmier (2014) *PNAS*[12] |

**Table S2. Benchmark data for simulations of DEER distance distributions (continued on following page).** Distances that could not be obtained due to steric hindrance are marked with *.

| | | Experimental | | | RosettaDEER | | MDDS | | MMM | | | | TagDock | | Pronox | | MtsslWizard | |
|---|---|---|---|---|---|---|---|---|---|---|---|---|---|---|---|---|---|---|
| | | | | | | | | | (ambient) | | (cryogenic) | | | | | | | |
| Res 1 | Res 2 | μ | σ | $C_\beta$-$C_\beta$ | μ | σ | μ | σ | μ | σ | μ | σ | μ | σ | μ | σ | μ | σ |
| **Bax (PDB: 1F16 H)** | | | | | | | | | | | | | | | | | | |
| 16 | 62 | 32.7 | 0.2 | 22.4 | 34.1 | 3.6 | 25.9 | 0.3 | 32.8 | 0.6 | 34.6 | 0.7 | 33.0 | 2.4 | 30.6 | 3.5 | 22.5 | 1.7 |
| 55 | 87 | 41.8 | 0.2 | 37.4 | 46.6 | 1.7 | 41.6 | 0.5 | 43.9 | 0.5 | 38.9 | 0.5 | 40.4 | 1.8 | 40.5 | 1.3 | 31.3 | 2.6 |
| 55 | 101 | 35.9 | 0.1 | 32.9 | 41.6 | 1.4 | 38.7 | 0.5 | 36.1 | 0.4 | 34.1 | 0.4 | 39.1 | 2.8 | 41.5 | 1.8 | 30.2 | 2.0 |
| 55 | 126 | 37.8 | 0.2 | 31.5 | 38.6 | 1.0 | 39.7 | 0.6 | 35.9 | 0.4 | 34.9 | 0.4 | 35.9 | 1.5 | 38.1 | 2.6 | 29.2 | 1.9 |
| 55 | 149 | 30.7 | 0.1 | 19.4 | 24.0 | 0.8 | 19.8 | 0.2 | 23.8 | 0.3 | 23.1 | 0.3 | 23.1 | 1.7 | 26.3 | 1.9 | 16.9 | 1.8 |
| 62 | 87 | 40.6 | 0.2 | 31.8 | 41.9 | 1.9 | 37.0 | 0.5 | 42.9 | 0.7 | 39.5 | 0.8 | 38.4 | 2.0 | 34.4 | 1.6 | 29.0 | 2.1 |
| 62 | 101 | 34.1 | 0.3 | 26.3 | 36.1 | 1.9 | 29.6 | 0.4 | 36.2 | 0.6 | 33.6 | 0.5 | 35.7 | 2.2 | 36.4 | 2.5 | 26.6 | 2.1 |
| 62 | 126 | 31.5 | 0.5 | 25.6 | 30.0 | 0.8 | 31.2 | 0.4 | 31.0 | 0.4 | 29.2 | 0.4 | 28.2 | 1.1 | 30.5 | 3.3 | 24.9 | 1.4 |
| 62 | 149 | 31.5 | 0.4 | 20.6 | 32.8 | 1.7 | 25.9 | 0.3 | 31.4 | 0.5 | 33.7 | 0.8 | 31.2 | 3.8 | 31.7 | 3.0 | 20.8 | 1.5 |
| 62 | 169 | 21.4 | 0.1 | 14.9 | 14.4 | 0.6 | 15.3 | 0.2 | 16.3 | 0.3 | 16.3 | 0.3 | 13.9 | 0.6 | 17.4 | 3.7 | 15.8 | 5.0 |
| 72 | 87 | 34.9 | 0.2 | 30.3 | 39.7 | 1.6 | 41.1 | 0.7 | 34.0 | 0.4 | 39.5 | 0.6 | 32.9 | 1.0 | 35.2 | 3.3 | 26.0 | 1.8 |
| 72 | 101 | 33.0 | 0.1 | 27.9 | 37.7 | 1.3 | 35.6 | 0.5 | 37.6 | 0.5 | 34.9 | 0.4 | 31.5 | 1.6 | 34.4 | 3.3 | 27.2 | 2.1 |
| 72 | 126 | 24.7 | 0.1 | 21.5 | 25.0 | 0.6 | 25.0 | 0.3 | 17.1 | 0.2 | 24.2 | 0.3 | 18.8 | 0.5 | 18.5 | 4.6 | 18.8 | 1.3 |
| 72 | 169 | 27.4 | 0.1 | 15.3 | 20.4 | 0.5 | 16.4 | 0.2 | 25.1 | 0.3 | 22.2 | 0.2 | 14.1 | 0.4 | 18.2 | 4.7 | 16.6 | 2.3 |
| 87 | 126 | 28.9 | 0.1 | 20.0 | 27.3 | 0.9 | 29.4 | 0.4 | 26.2 | 0.3 | 25.9 | 0.3 | 20.2 | 0.8 | 22.6 | 3.3 | 16.4 | 2.5 |
| 101 | 126 | 34.0 | 0.0 | 32.2 | 38.9 | 1.0 | 37.5 | 0.4 | 37.1 | 0.4 | 39.1 | 0.5 | 32.1 | 1.3 | 34.1 | 3.7 | 31.0 | 2.2 |
| 101 | 149 | 29.1 | 0.1 | 19.3 | 28.4 | 1.5 | 29.5 | 0.5 | 21.8 | 0.2 | 23.4 | 0.4 | 28.1 | 2.5 | 30.2 | 2.4 | 19.9 | 1.1 |
| 101 | 169 | 26.2 | 0.1 | 26.2 | 36.1 | 1.1 | 34.0 | 0.4 | 38.5 | 0.5 | 30.6 | 0.3 | 32.7 | 1.3 | 34.9 | 4.3 | 29.1 | 2.3 |
| 126 | 169 | 39.8 | 0.1 | 34.3 | 40.3 | 0.9 | 39.5 | 0.5 | 38.0 | 0.4 | 39.3 | 0.5 | 30.7 | 0.9 | 33.5 | 3.8 | 31.7 | 1.9 |
| 72 | 186 | 29.7 | 0.8 | 25.8 | 25.6 | 0.8 | 25.2 | 0.3 | 23.6 | 0.3 | 25.1 | 0.1 | 24.8 | 0.8 | 24.0 | 3.2 | 19.1 | 1.3 |
| 87 | 186 | 27.0 | 1.3 | 25.1 | 27.2 | 1.0 | 30.1 | 0.5 | 29.0 | 0.4 | 24.6 | 0.0 | 28.9 | 1.3 | 31.9 | 3.1 | 19.6 | 1.3 |
| **Mhp1 (PDB: 2JLN)** | | | | | | | | | | | | | | | | | | |
| 30 | 163 | 32.5 | 0.0 | 17.5 | 22.7 | 0.7 | 22.5 | 0.3 | 23.6 | 0.3 | 21.2 | 0.3 | 25.5 | 1.2 | 22.1 | 3.1 | 17.7 | 1.5 |
| 30 | 243 | 25.6 | 0.1 | 22.9 | 25.6 | 0.6 | 27.8 | 0.3 | 26.3 | 0.3 | 24.8 | 0.3 | 19.8 | 0.9 | 28.4 | 3.5 | 23.9 | 1.1 |
| 30 | 338 | 45.8 | 0.5 | 32.3 | 38.5 | 1.2 | 39.5 | 0.6 | 40.5 | 0.5 | 38.9 | 0.5 | 41.3 | 2.1 | 41.6 | 2.6 | 32.2 | 1.9 |
| 51 | 278 | 24.1 | 0.2 | 14.3 | 18.6 | 0.6 | 18.7 | 0.3 | 21.4 | 0.2 | 19.2 | 0.2 | 23.4 | 0.8 | 21.1 | 4.1 | 16.3 | 2.6 |
| 63 | 285 | 34.7 | 0.0 | 25.9 | 32.4 | 1.3 | 35.6 | 0.6 | 35.8 | 0.5 | 33.8 | 0.4 | 33.4 | 1.7 | 36.5 | 2.3 | 26.3 | 1.9 |
| 63 | 362 | 32.6 | 0.0 | 29.4 | 35.1 | 1.2 | 36.0 | 0.6 | 38.4 | 4.8 | 35.1 | 0.5 | 36.5 | 1.4 | 38.7 | 2.5 | 29.8 | 1.7 |
| 136 | 278 | 27.0 | 0.0 | 25.9 | 26.9 | 1.1 | 32.3 | 0.4 | 31.4 | 0.4 | 31.2 | 0.4 | 31.2 | 1.1 | 31.2 | 2.6 | 25.8 | 1.8 |
| 136 | 349 | 33.4 | 0.1 | 30.3 | 34.2 | 1.0 | 36.6 | 0.5 | 34.3 | 0.3 | 32.2 | 0.3 | 31.3 | 0.9 | 36.6 | 4.5 | 30.4 | 2.0 |
| 144 | 278 | 33.7 | 0.1 | 31.6 | 36.0 | 1.1 | 39.0 | 0.5 | 44.6 | 0.7 | 38.1 | 0.4 | 39.5 | 1.7 | 42.4 | 2.1 | 32.5 | 2.1 |
| 159 | 324 | 24.0 | 0.1 | 15.0 | 17.6 | 1.0 | 15.3 | 0.2 | 13.8 | 0.2 | 13.8 | 0.3 | 16.1 | 0.6 | 14.9 | 2.2 | 17.0 | 1.6 |
| 163 | 243 | 45.9 | 0.0 | 34.8 | 40.1 | 1.6 | 42.4 | 0.6 | 42.3 | 0.6 | 39.9 | 0.6 | 40.7 | 2.1 | 43.6 | 2.2 | 35.2 | 1.7 |
| 184 | 278 | 27.0 | 0.0 | 16.4 | 20.5 | 0.9 | 25.5 | 0.3 | 20.7 | 0.2 | 22.5 | 0.4 | 23.2 | 1.0 | 24.5 | 2.6 | 24.8 | 0.9 |
| 234 | 338 | 42.6 | 0.0 | 25.7 | 31.1 | 1.3 | 25.9 | 0.5 | 30.3 | 0.5 | 29.7 | 0.6 | 32.0 | 1.1 | 32.9 | 3.9 | n/a* | n/a* |
| 243 | 338 | 58.7 | 0.2 | 41.8 | 47.5 | 1.7 | 49.5 | 0.7 | 49.1 | 0.5 | 47.9 | 0.8 | 49.3 | 2.3 | 32.9 | 3.9 | 41.7 | 2.3 |
| 278 | 349 | 36.6 | 0.0 | 36.9 | 40.9 | 1.0 | 40.9 | 0.5 | 46.0 | 0.5 | 38.9 | 0.4 | 43.7 | 1.8 | 45.1 | 3.4 | 37.2 | 2.1 |
| 278 | 362 | 33.9 | 0.1 | 30.5 | 36.6 | 1.1 | 36.8 | 0.6 | 42.7 | 0.6 | 35.9 | 0.4 | 39.7 | 1.7 | 40.2 | 3.0 | 30.5 | 1.7 |
| 285 | 349 | 33.1 | 0.0 | 31.0 | 32.9 | 0.8 | 31.3 | 0.4 | 31.7 | 0.3 | 31.1 | 0.3 | 32.1 | 1.0 | 31.9 | 3.7 | 31.0 | 1.5 |
| 349 | 362 | 31.4 | 0.0 | 19.1 | 23.7 | 0.6 | 23.6 | 0.3 | 25.7 | 0.3 | 21.2 | 0.2 | 24.3 | 0.8 | 24.6 | 3.9 | 21.1 | 1.2 |
| **ExoU (PDB: 3TU3)** | | | | | | | | | | | | | | | | | | |
| 592 | 636 | 26.8 | 0.1 | 26.2 | 27.7 | 0.7 | 21.9 | 0.3 | 25.2 | 0.3 | 22.4 | 0.3 | 19.3 | 0.9 | 26.0 | 3.8 | 25.6 | 1.5 |
| 592 | 649 | 24.7 | 0.0 | 24.8 | 26.1 | 0.7 | 25.5 | 0.3 | 25.3 | 0.2 | 24.4 | 0.3 | 20.3 | 1.0 | 26.4 | 4.7 | 25.7 | 1.6 |
| 598 | 680 | 27.8 | 0.1 | 21.5 | 20.7 | 0.6 | 21.5 | 0.3 | 25.6 | 0.3 | 27.3 | 0.3 | 22.4 | 0.7 | 26.7 | 2.7 | 22.0 | 1.4 |
| 629 | 645 | 25.1 | 0.1 | 23.5 | 29.8 | 0.8 | 27.9 | 0.3 | 27.7 | 0.3 | 27.9 | 0.3 | 22.4 | 0.7 | 26.7 | 3.7 | 21.6 | 1.4 |
| 636 | 645 | 23.4 | 0.1 | 16.2 | 24.7 | 0.9 | 24.7 | 0.3 | 24.0 | 0.3 | 21.5 | 0.2 | 22.0 | 0.7 | 20.2 | 4.1 | 16.9 | 1.9 |
| 636 | 649 | 22.0 | 0.1 | 11.9 | 20.0 | 0.6 | 20.5 | 0.3 | 20.7 | 0.2 | 19.1 | 0.3 | 16.3 | 0.4 | 14.5 | 4.3 | 15.8 | 4.7 |
| 636 | 657 | 23.5 | 0.1 | 11.9 | 19.8 | 0.6 | 20.3 | 0.3 | 19.6 | 0.2 | 29.3 | 0.5 | 19.0 | 0.7 | 15.8 | 2.6 | 16.6 | 2.3 |
| 636 | 672 | 29.4 | 0.2 | 23.9 | 31.1 | 1.2 | 32.2 | 0.5 | 30.1 | 0.5 | 31.4 | 0.4 | 28.2 | 1.5 | 28.4 | 1.4 | 21.1 | 1.5 |
| 636 | 677 | 27.9 | 0.2 | 18.9 | 28.3 | 1.1 | 27.3 | 0.4 | 28.0 | 0.4 | 27.5 | 0.4 | 26.1 | 1.5 | 28.7 | 2.1 | 18.4 | 1.3 |
| 636 | 682 | 25.4 | 0.1 | 18.4 | 25.9 | 1.0 | 24.6 | 0.4 | 25.3 | 0.3 | 24.9 | 0.3 | 25.9 | 1.1 | 24.9 | 2.4 | 19.0 | 1.1 |
| 649 | 672 | 26.5 | 0.2 | 20.7 | 24.1 | 0.6 | 21.4 | 0.2 | 22.3 | 0.3 | 23.1 | 0.3 | 19.6 | 0.8 | 21.8 | 3.3 | 18.1 | 1.5 |

| | | Experimental | | | RosettaDEER | | MDDS | | MMM (ambient) | | MMM (cryogenic) | | TagDock | | Pronox | | MtsslWizard | |
|---|---|---|---|---|---|---|---|---|---|---|---|---|---|---|---|---|---|---|
| Res 1 | Res 2 | μ | σ | Cβ-Cβ | μ | σ | μ | σ | μ | σ | μ | σ | μ | σ | μ | σ | μ | σ |
| **CDB3 (PDB: 1HYN R/S)** | | | | | | | | | | | | | | | | | | |
| 84 | 84 | 31.5 | 0.1 | 31.0 | 30.4 | 1.1 | 21.7 | 0.3 | 25.5 | 0.3 | 20.6 | 0.4 | 32.7 | 1.6 | 32.1 | 1.7 | 30.9 | 2.1 |
| 96 | 96 | 31.2 | 0.1 | 33.9 | 36.4 | 0.9 | 31.5 | 0.4 | 32.5 | 0.4 | 32.2 | 0.3 | 30.2 | 1.1 | 26.8 | 2.7 | 31.9 | 1.7 |
| 105 | 105 | 13.6 | 0.0 | 14.9 | 15.7 | 0.6 | 14.5 | 0.2 | 23.4 | 0.4 | 20.4 | 0.3 | 11.0 | 0.7 | 22.4 | 4.5 | 15.7 | 5.4 |
| 116 | 116 | 11.1 | 0.1 | 23.9 | 22.5 | 0.5 | 22.0 | 0.3 | 22.4 | 0.2 | 23.7 | 0.3 | 24.5 | 1.1 | 20.8 | 3.8 | 23.5 | 1.2 |
| 142 | 142 | 28.5 | 0.1 | 37.8 | 34.1 | 0.7 | 34.4 | 0.4 | 37.3 | 0.4 | 32.6 | 0.5 | 44.5 | 4.8 | 33.8 | 2.8 | 38.3 | 2.7 |
| 199 | 199 | 31.8 | 0.1 | 29.2 | 32.5 | 1.8 | 30.7 | 0.4 | 27.1 | 0.4 | 36.2 | 0.5 | 22.0 | 0.7 | 35.0 | 0.7 | 30.1 | 1.8 |
| 208 | 208 | 39.2 | 0.1 | 44.0 | 46.5 | 1.1 | 37.5 | 0.6 | 38.9 | 0.4 | 40.3 | 0.4 | 36.1 | 2.9 | 43.5 | 5.2 | 44.2 | 4.0 |
| 277 | 277 | 30.6 | 0.1 | 27.9 | 32.6 | 0.9 | 36.0 | 0.5 | 35.3 | 0.4 | 37.4 | 0.5 | 31.2 | 1.2 | 27.5 | 1.2 | 29.0 | 2.7 |
| 290 | 290 | 26.2 | 0.2 | 33.3 | 38.4 | 0.9 | 40.3 | 0.5 | 21.5 | 0.4 | 19.8 | 0.5 | 26.3 | 1.2 | 22.1 | 1.6 | 33.9 | 6.0 |
| 312 | 312 | 28.7 | 0.0 | 24.4 | 25.8 | 0.9 | 21.5 | 0.2 | 25.2 | 0.3 | 26.4 | 0.4 | 20.1 | 0.6 | 30.6 | 2.8 | 24.1 | 2.1 |
| 340 | 340 | 21.6 | 0.1 | 24.3 | 30.6 | 0.8 | 30.8 | 0.4 | 29.9 | 0.4 | 31.9 | 0.5 | 25.9 | 0.8 | 30.9 | 2.9 | 23.9 | 1.9 |
| 342 | 342 | 12.6 | 0.1 | 20.2 | 21.1 | 0.8 | 25.4 | 0.3 | 23.2 | 0.4 | 23.8 | 0.7 | 30.1 | 3.6 | 18.6 | 2.7 | 21.9 | 1.3 |
| 343 | 343 | 16.9 | 0.2 | 25.1 | 32.1 | 1.4 | 31.4 | 0.5 | 31.3 | 0.4 | 32.4 | 0.5 | 23.6 | 0.9 | 34.9 | 2.2 | 26.7 | 2.2 |
| 344 | 344 | 40.5 | 0.2 | 33.1 | 40.7 | 1.1 | 43.1 | 0.6 | 39.4 | 0.4 | 38.5 | 0.5 | 42.0 | 3.4 | 40.5 | 2.8 | 35.7 | 1.8 |
| 345 | 345 | 26.7 | 0.1 | 30.3 | 36.6 | 1.1 | 37.7 | 0.5 | 38.9 | 0.5 | 36.9 | 0.4 | 26.3 | 1.2 | 35.6 | 4.6 | 31.4 | 2.9 |
| **Rhodopsin (PDB: 1GZM A)** | | | | | | | | | | | | | | | | | | |
| 63 | 241 | 34.4 | 0.2 | 26.9 | 40.6 | 1.6 | 39.1 | 0.5 | 37.8 | 0.4 | 38.8 | 0.5 | 36.4 | 1.5 | 42.4 | 3.2 | 31.0 | 2.3 |
| 63 | 252 | 31.1 | 0.1 | 25.8 | 36.3 | 1.7 | 35.3 | 0.5 | 33.1 | 0.4 | 34.0 | 0.5 | 31.6 | 1.5 | 35.7 | 2.5 | 26.0 | 1.8 |
| 63 | 326 | 26.3 | 0.1 | 17.9 | 26.3 | 1.4 | 24.2 | 0.4 | 22.4 | 0.4 | 22.7 | 0.4 | 24.2 | 1.3 | 29.2 | 2.7 | 18.5 | 1.4 |
| 74 | 137 | 22.3 | 0.1 | 14.3 | 21.1 | 2.0 | 19.7 | 0.3 | 22.9 | 0.3 | 23.1 | 0.4 | 19.5 | 1.3 | 21.0 | 0.6 | 16.6 | 2.2 |
| 74 | 225 | 31.4 | 0.2 | 21.4 | 33.7 | 2.1 | 30.5 | 0.4 | 33.0 | 0.5 | 34.1 | 0.6 | 31.3 | 2.0 | 36.0 | 0.7 | 24.1 | 1.7 |
| 74 | 252 | 29.2 | 0.1 | 18.9 | 31.0 | 2.0 | 30.8 | 0.6 | 30.9 | 0.5 | 29.4 | 0.6 | 29.6 | 2.8 | 32.0 | 1.5 | 20.9 | 1.0 |
| 74 | 308 | 27.8 | 0.4 | 19.3 | 28.6 | 1.8 | 29.6 | 0.5 | 30.1 | 0.6 | 28.9 | 0.6 | 26.5 | 1.6 | 29.5 | 1.6 | 20.4 | 0.9 |
| 137 | 326 | 44.0 | 0.3 | 32.1 | 37.7 | 2.6 | 37.1 | 0.5 | 35.9 | 0.7 | 36.0 | 0.8 | 34.7 | 3.4 | 35.8 | 4.9 | 32.9 | 2.3 |
| 151 | 241 | 32.5 | 0.0 | 29.7 | 37.0 | 1.3 | 38.4 | 0.6 | 37.2 | 0.5 | 35.5 | 0.5 | 33.5 | 1.4 | 37.5 | 3.5 | 28.0 | 2.9 |
| 151 | 308 | 35.9 | 0.1 | 29.3 | 37.3 | 1.6 | 39.6 | 0.6 | 38.8 | 0.5 | 36.7 | 0.6 | 35.3 | 1.6 | 38.5 | 2.6 | 28.8 | 1.7 |
| 151 | 326 | 31.8 | 0.1 | 31.0 | 35.1 | 1.1 | 36.0 | 0.5 | 30.0 | 0.4 | 31.2 | 0.4 | 34.2 | 1.2 | 37.5 | 3.7 | 31.2 | 2.2 |
| 225 | 252 | 25.0 | 0.2 | 16.0 | 21.8 | 0.8 | 24.2 | 0.3 | 20.5 | 0.2 | 21.1 | 0.3 | 24.5 | 1.1 | 25.2 | 3.4 | 18.1 | 1.3 |
| 225 | 308 | 34.0 | 0.2 | 11.5 | 35.2 | 1.5 | 36.5 | 0.5 | 36.5 | 0.5 | 34.0 | 0.6 | 32.9 | 1.1 | 36.6 | 2.7 | 27.7 | 1.4 |
| 252 | 326 | 32.7 | 0.2 | 23.3 | 26.1 | 1.1 | 25.4 | 0.3 | 28.2 | 0.4 | 26.0 | 0.3 | 21.2 | 0.7 | 23.9 | 4.7 | 23.9 | 1.8 |
| Correlation | | | | 0.42 | 0.56 | 0.02 | 0.48 | 0.01 | 0.52 | 0.00 | 0.51 | 0.01 | 0.54 | 0.01 | 0.43 | 0.01 | 0.43 | 0.02 |

**Table S3. List of spin-labeled proteins in the protein data bank.**

| PDB | Beta carbon | | | Nitroxide nitrogen | | | Nitroxide oxygen | | | Angle | Relative Distance |
|---|---|---|---|---|---|---|---|---|---|---|---|
| | x | y | z | x | y | z | x | y | z | | |
| 1ZYT (A) | 45.34 | -7.24 | 7.94 | 45.93 | -3.81 | 14.56 | 46.54 | -3.29 | 15.49 | 5.23 | 0.83 |
| 2CUU (A) | 31.74 | 9.11 | -11.29 | 37.09 | 11.32 | -15.90 | 38.33 | 11.54 | -16.04 | 7.24 | 0.84 |
| 2CUU (A) | 31.74 | 9.11 | -11.29 | 34.84 | 8.96 | -16.89 | 35.44 | 8.64 | -17.97 | 3.39 | 0.78 |
| 2IGC (A) | 0.63 | 23.03 | 97.36 | -1.83 | 18.50 | 98.24 | -2.70 | 18.07 | 98.95 | 11.49 | 0.81 |
| 2NTH (A) | -11.76 | 35.89 | 63.20 | -6.93 | 35.75 | 67.89 | -6.25 | 35.86 | 68.89 | 3.06 | 0.80 |
| 2OU8 (A) | 0.09 | 23.50 | 99.06 | -4.09 | 18.69 | 97.25 | -5.21 | 18.25 | 97.24 | 7.13 | 0.82 |
| 2OU8 (A) | 0.09 | 23.50 | 99.06 | 3.34 | 18.79 | 103.07 | 4.14 | 17.91 | 102.86 | 9.08 | 0.86 |
| 2OU9 (A) | 20.05 | 11.78 | 46.38 | 18.14 | 7.36 | 46.35 | 18.37 | 6.51 | 45.54 | 15.13 | 0.84 |
| 2Q9D (A) | 25.85 | 29.50 | 8.70 | 28.07 | 33.43 | 6.03 | 27.97 | 34.62 | 5.86 | 10.99 | 0.80 |
| 2Q9E (A) | 22.82 | 4.02 | 77.45 | 22.72 | 8.29 | 80.31 | 22.16 | 9.30 | 79.97 | 14.55 | 0.86 |
| 2Q9E (B) | 16.24 | -6.49 | 118.33 | 20.24 | -6.61 | 121.49 | 21.32 | -6.13 | 121.42 | 13.03 | 0.83 |
| 2Q9E (C) | -13.47 | -10.74 | 116.70 | -14.18 | -16.97 | 119.08 | -14.98 | -17.70 | 119.55 | 8.08 | 0.84 |
| 2XGA (A) | 27.64 | 36.01 | -12.25 | 33.29 | 35.15 | -9.09 | 34.17 | 35.48 | -8.27 | 6.37 | 0.80 |
| 2XGA (A) | 53.10 | 12.38 | 32.02 | 53.37 | 13.62 | 25.61 | 53.22 | 13.41 | 24.38 | 5.53 | 0.80 |
| 2XIU (A) | 30.89 | 11.72 | 13.12 | 32.52 | 18.09 | 14.29 | 32.68 | 19.22 | 13.87 | 7.07 | 0.82 |
| 2XIU (A) | 31.18 | 11.32 | 12.96 | 12.88 | 1.28 | 14.75 | 12.82 | 0.14 | 15.17 | 3.59 | 0.96 |
| 3G3V (A) | 32.01 | 9.42 | -11.26 | 37.17 | 11.36 | -16.07 | 38.43 | 11.38 | -16.25 | 7.83 | 0.84 |
| 3G3V (A) | 32.01 | 9.42 | -11.26 | 34.94 | 8.76 | -16.91 | 35.47 | 8.19 | -17.91 | 5.30 | 0.79 |
| 3M8B (A) | -36.14 | -15.28 | 3.58 | -34.89 | -19.33 | 7.52 | -34.61 | -20.63 | 7.42 | 12.64 | 0.82 |
| 3M8D (A) | 30.56 | -24.22 | -34.24 | 33.53 | -21.66 | -29.05 | 34.71 | -20.81 | -28.85 | 11.98 | 0.82 |
| 3MPN (A) | 46.41 | 34.94 | 18.10 | 47.66 | 35.80 | 25.63 | 47.41 | 35.92 | 27.00 | 4.61 | 0.80 |
| 3MPQ (A) | 29.69 | 31.87 | -0.63 | 29.42 | 28.90 | -4.58 | 29.24 | 27.53 | -4.75 | 15.33 | 0.79 |
| 3RGM (A) | 28.45 | -2.69 | -21.73 | 34.19 | 1.12 | -20.61 | 34.77 | 2.03 | -19.73 | 9.04 | 0.82 |
| 3RGN (A) | -34.31 | 44.87 | -34.41 | -31.48 | 49.85 | -38.26 | -31.67 | 51.20 | -38.45 | 10.25 | 0.83 |
| 3STZ (C) | 151.02 | 114.96 | -17.24 | 156.25 | 149.05 | -22.46 | 156.30 | 149.33 | -21.26 | 2.51 | 1.00 |
| 3STZ (C) | 155.90 | 146.71 | -29.68 | 156.25 | 149.05 | -22.46 | 156.30 | 149.33 | -21.26 | 1.10 | 0.81 |
| 3V3X (A) | 8.06 | 35.66 | 44.62 | 7.17 | 40.73 | 39.31 | 6.94 | 41.37 | 38.29 | 2.62 | 0.81 |
| 3V3X (B) | 20.59 | 27.24 | 17.69 | 13.12 | 27.74 | 16.14 | 12.09 | 27.49 | 15.56 | 4.78 | 0.83 |
| 3V3X (B) | 20.43 | 27.14 | 17.71 | 14.29 | 28.00 | 16.08 | 13.28 | 27.42 | 15.67 | 8.53 | 0.82 |
| 3V3X (B) | 20.826 | 30.531 | -11.28 | 26.72 | 40.34 | 48.40 | 27.59 | 40.28 | 47.56 | 1.13 | 1.02 |
| 3V3X (C) | 5.82 | 27.48 | 36.57 | 9.99 | 23.00 | 39.69 | 10.62 | 22.08 | 40.20 | 2.00 | 0.80 |
| 3V3X (D) | 22.506 | 23.768 | 6.859 | 27.85 | 21.68 | 8.62 | 28.91 | 21.03 | 8.59 | 5.65 | 0.78 |
| 3V3X (D) | 22.401 | 24.15 | 6.799 | 28.74 | 23.01 | 9.39 | 29.87 | 22.71 | 9.77 | 1.39 | 0.80 |
| 3V3X (D) | 28.74 | 38.19 | 32.17 | 31.88 | 41.91 | 26.95 | 32.61 | 42.58 | 26.22 | 2.71 | 0.81 |
| 4EK1 (A) | 28.41 | -27.82 | 0.47 | 29.53 | -29.48 | 7.15 | 29.96 | -30.24 | 8.29 | 5.64 | 0.78 |
| 4EK1 (B) | 17.67 | -27.74 | 35.80 | 19.44 | -29.86 | 41.92 | 19.43 | -30.90 | 42.84 | 8.46 | 0.80 |
| 5BMG (A) | 68.09 | 8.92 | 26.62 | 69.95 | 14.65 | 20.03 | 69.36 | 15.35 | 21.21 | 10.97 | 1.08 |
| 5BMG (B) | 70.46 | 8.41 | 30.86 | 68.76 | 14.18 | 32.79 | 68.81 | 15.31 | 33.24 | 4.57 | 0.79 |
| 5BMG (B) | 70.46 | 8.41 | 30.86 | 74.05 | 10.31 | 26.90 | 74.00 | 9.10 | 27.04 | 13.52 | 1.13 |
| 5BMG (D) | 64.35 | -31.08 | 48.92 | 66.58 | -25.65 | 51.29 | 67.33 | -24.68 | 51.37 | 5.89 | 0.80 |
| 5BMG (E) | 61.98 | 23.34 | 48.91 | 61.65 | -28.81 | 44.96 | 62.20 | -29.35 | 44.01 | 1.45 | 0.99 |
| 5BMG (F) | 59.71 | -22.93 | 53.48 | 62.05 | -28.84 | 56.84 | 61.78 | -29.62 | 57.73 | 7.70 | 0.84 |
| 5BMG (G) | 67.80 | 17.05 | 26.41 | 63.62 | 11.03 | 27.98 | 63.33 | 10.00 | 27.39 | 8.47 | 0.87 |
| 5BMH (A) | 0.05 | -10.04 | -13.40 | -1.59 | -8.86 | -18.40 | -1.45 | -8.90 | -19.62 | 8.29 | 0.77 |
| 5BMH (A) | 0.21 | -10.02 | -13.65 | -1.66 | -8.67 | -18.34 | -1.55 | -8.70 | -19.56 | 9.38 | 0.77 |
| 5BMI (A) | -12.45 | 12.28 | -1.86 | -7.06 | 15.58 | -4.29 | -6.04 | 15.52 | -4.96 | 7.42 | 0.83 |
| 5I26 (A) | 94.17 | 13.89 | 136.21 | 97.47 | 15.70 | 141.52 | 97.75 | 16.31 | 142.46 | 4.80 | 0.81 |
| 5I26 (B) | 121.29 | 25.02 | 121.83 | 121.69 | 18.58 | 121.08 | 122.34 | 17.62 | 121.18 | 0.55 | 0.99 |
| 5I26 (C) | 90.81 | 62.80 | 115.49 | 85.28 | 65.25 | 117.86 | 84.85 | 65.22 | 118.94 | 0.92 | 1.01 |
| 5I26 (D) | 75.61 | 38.89 | 96.79 | 78.64 | 41.52 | 91.87 | 78.33 | 42.28 | 91.05 | 0.89 | 1.00 |
| 5I28 (A) | 253.06 | 53.98 | 41.53 | 256.95 | 56.60 | 46.87 | 257.07 | 57.16 | 47.86 | 0.50 | 1.00 |
| 5I28 (B) | 241.07 | 51.61 | 72.62 | 237.23 | 53.97 | 67.15 | 237.07 | 54.42 | 66.10 | 0.71 | 1.00 |
| 5I28 (C) | 254.91 | 59.75 | 42.24 | 248.27 | 57.34 | 42.90 | 247.35 | 56.89 | 43.43 | 0.32 | 1.01 |
| 5I28 (D) | 261.00 | 7.94 | 25.94 | 260.59 | 4.56 | 22.84 | 259.61 | 4.31 | 22.27 | 0.25 | 1.01 |
| 5I28 (E) | 242.01 | 18.87 | 3.05 | 238.04 | 15.54 | -1.77 | 237.84 | 14.60 | -2.40 | 4.49 | 0.96 |
| 5I28 (F) | 290.39 | 18.69 | 26.81 | 285.83 | 18.41 | 20.66 | 285.50 | 18.81 | 19.63 | 1.81 | 1.01 |
| 5I28 (G) | 223.92 | 68.30 | 42.05 | 222.89 | 64.06 | 47.69 | 222.43 | 63.01 | 47.87 | 2.15 | 0.98 |
| 5I28 (H) | 268.83 | 69.81 | 68.41 | 271.91 | 65.90 | 65.16 | 272.94 | 65.38 | 65.17 | 2.10 | 0.89 |
| 5I28 (I) | 270.04 | 41.64 | 65.86 | 269.40 | 45.76 | 70.37 | 269.61 | 46.73 | 70.97 | 5.44 | 0.80 |
| 5I28 (J) | 222.99 | 43.38 | 41.56 | 219.19 | 47.62 | 46.17 | 219.27 | 48.46 | 46.97 | 6.80 | 0.85 |
| 5JDT (A) | 34.90 | -28.61 | -16.81 | 32.74 | -24.46 | -11.81 | 32.18 | -23.58 | -11.01 | 2.52 | 0.78 |
| 5JDT (A) | 35.02 | -28.35 | -16.57 | 32.87 | -24.65 | -11.87 | 32.37 | -23.92 | -10.88 | 0.85 | 0.76 |

**Table S4. Experimental DEER decay parameters (continued on following page).**

| Res 1 | Res 2 | k | λ | Duration | |
|---|---|---|---|---|---|
| | | | | Oscillations | μs |
| Bax (PDB: 1F16 H) | | | | | |
| 16 | 62 | -0.01 | 0.389 | 4.45 | 3.00 |
| 55 | 87 | -0.01 | 0.348 | 2.64 | 3.70 |
| 55 | 101 | 0.00 | 0.342 | 3.59 | 3.20 |
| 55 | 126 | 0.00 | 0.337 | 2.99 | 3.10 |
| 55 | 149 | -0.01 | 0.286 | 4.85 | 2.70 |
| 62 | 87 | -0.01 | 0.389 | 2.71 | 3.50 |
| 62 | 101 | 0.00 | 0.397 | 3.54 | 2.70 |
| 62 | 126 | -0.27 | 0.55 | 7.66 | 4.62 |
| 62 | 149 | -0.01 | 0.442 | 4.49 | 2.70 |
| 62 | 169 | -0.01 | 0.286 | 12.73 | 2.40 |
| 72 | 87 | -0.01 | 0.388 | 3.93 | 3.20 |
| 72 | 101 | -0.01 | 0.304 | 4.77 | 3.30 |
| 72 | 126 | 0.00 | 0.237 | 8.62 | 2.50 |
| 72 | 169 | -0.01 | 0.382 | 7.55 | 3.00 |
| 87 | 126 | -0.01 | 0.234 | 4.33 | 2.00 |
| 101 | 126 | 0.00 | 0.195 | 2.26 | 1.70 |
| 101 | 149 | -0.01 | 0.206 | 3.18 | 1.50 |
| 101 | 169 | -0.01 | 0.319 | 5.23 | 1.80 |
| 126 | 169 | 0.00 | 0.356 | 2.47 | 3.00 |
| 72 | 186 | -0.01 | 0.135 | 3.38 | 1.70 |
| 87 | 186 | -0.01 | 0.175 | 4.48 | 1.70 |
| Mhp1 (PDB: 2JLN) | | | | | |
| 30 | 163 | -0.05 | 0.115 | 4.47 | 2.95 |
| 30 | 243 | -0.08 | 0.127 | 6.32 | 2.05 |
| 30 | 338 | 0.00 | 0.125 | 1.16 | 2.14 |
| 51 | 278 | -0.05 | 0.171 | 6.48 | 1.74 |
| 63 | 285 | -0.05 | 0.072 | 3.66 | 2.94 |
| 63 | 362 | -0.04 | 0.076 | 3.82 | 2.54 |
| 136 | 278 | -0.04 | 0.099 | 6.42 | 2.44 |
| 136 | 349 | -0.04 | 0.064 | 4.10 | 2.94 |
| 144 | 278 | -0.03 | 0.13 | 3.72 | 2.74 |
| 159 | 324 | -0.12 | 0.218 | 7.72 | 2.05 |
| 163 | 243 | -0.02 | 0.1 | 1.74 | 3.24 |
| 184 | 278 | -0.07 | 0.163 | 6.22 | 2.34 |
| 234 | 338 | -0.03 | 0.064 | 1.78 | 2.65 |
| 243 | 338 | -0.03 | 0.252 | 0.89 | 3.45 |
| 278 | 349 | -0.05 | 0.052 | 3.13 | 2.94 |
| 278 | 362 | -0.04 | 0.143 | 3.65 | 2.74 |
| 285 | 349 | -0.02 | 0.057 | 3.93 | 2.74 |
| 349 | 362 | -0.04 | 0.117 | 4.62 | 2.74 |
| ExoU (PDB: 3TU3) | | | | | |
| 592 | 636 | -0.05 | 0.127 | 7.07 | 2.62 |
| 592 | 649 | -0.01 | 0.081 | 10.05 | 2.91 |
| 598 | 680 | -0.05 | 0.096 | 7.12 | 2.93 |
| 629 | 645 | -0.01 | 0.088 | 6.66 | 2.02 |
| 636 | 645 | -0.02 | 0.117 | 11.10 | 2.72 |
| 636 | 649 | -0.06 | 0.123 | 12.74 | 2.62 |
| 636 | 657 | -0.03 | 0.115 | 10.57 | 2.62 |
| 636 | 672 | -0.06 | 0.149 | 5.36 | 2.62 |
| 636 | 677 | -0.03 | 0.18 | 5.85 | 2.43 |
| 636 | 682 | -0.03 | 0.164 | 8.71 | 2.74 |
| 649 | 672 | -0.04 | 0.2 | 6.53 | 2.34 |

| Res 1 | Res 2 | k | λ | Duration | |
|---|---|---|---|---|---|
| | | | | Oscillations | µs |
| CDB3 (PDB: 1HYN R/S) | | | | | |
| 84 | 84 | -0.06 | 0.185 | 2.65 | 1.59 |
| 96 | 96 | -0.07 | 0.177 | 2.72 | 1.59 |
| 105 | 105 | -0.05 | 0.122 | 16.35 | 0.78 |
| 116 | 116 | -0.05 | 0.108 | 29.81 | 0.78 |
| 142 | 142 | -0.03 | 0.071 | 3.58 | 1.59 |
| 199 | 199 | -0.10 | 0.146 | 3.86 | 2.38 |
| 208 | 208 | -0.06 | 0.062 | 2.58 | 2.98 |
| 277 | 277 | -0.03 | 0.19 | 3.60 | 1.99 |
| 290 | 290 | -0.08 | 0.195 | 12.88 | 4.46 |
| 312 | 312 | -0.03 | 0.078 | 4.36 | 1.99 |
| 340 | 340 | -0.04 | 0.243 | 12.81 | 2.47 |
| 342 | 342 | -0.06 | 0.163 | 40.76 | 1.57 |
| 343 | 343 | -0.08 | 0.17 | 17.06 | 1.59 |
| 344 | 344 | -0.07 | 0.162 | 2.37 | 3.02 |
| 345 | 345 | -0.04 | 0.228 | 8.13 | 2.98 |
| Rhodopsin (PDB: 1GZM A) | | | | | |
| 63 | 241 | -0.36 | 0.326 | 3.26 | 2.56 |
| 63 | 252 | -0.05 | 0.233 | 3.42 | 1.97 |
| 63 | 326 | -0.09 | 0.101 | 4.17 | 1.46 |
| 74 | 137 | -0.16 | 0.218 | 8.29 | 1.77 |
| 74 | 225 | -0.13 | 0.234 | 2.96 | 1.77 |
| 74 | 252 | -0.10 | 0.214 | 3.70 | 1.77 |
| 74 | 308 | -0.14 | 0.413 | 4.77 | 1.98 |
| 137 | 326 | 0.02 | 0.25 | 1.55 | 2.54 |
| 151 | 241 | -0.18 | 0.085 | 3.84 | 2.54 |
| 151 | 308 | -0.03 | 0.133 | 2.21 | 1.97 |
| 151 | 326 | -0.05 | 0.146 | 4.01 | 2.47 |
| 225 | 252 | -0.35 | 0.277 | 3.90 | 1.17 |
| 225 | 308 | -0.25 | 0.292 | 3.28 | 2.47 |
| 252 | 326 | -0.04 | 0.282 | 2.92 | 1.97 |

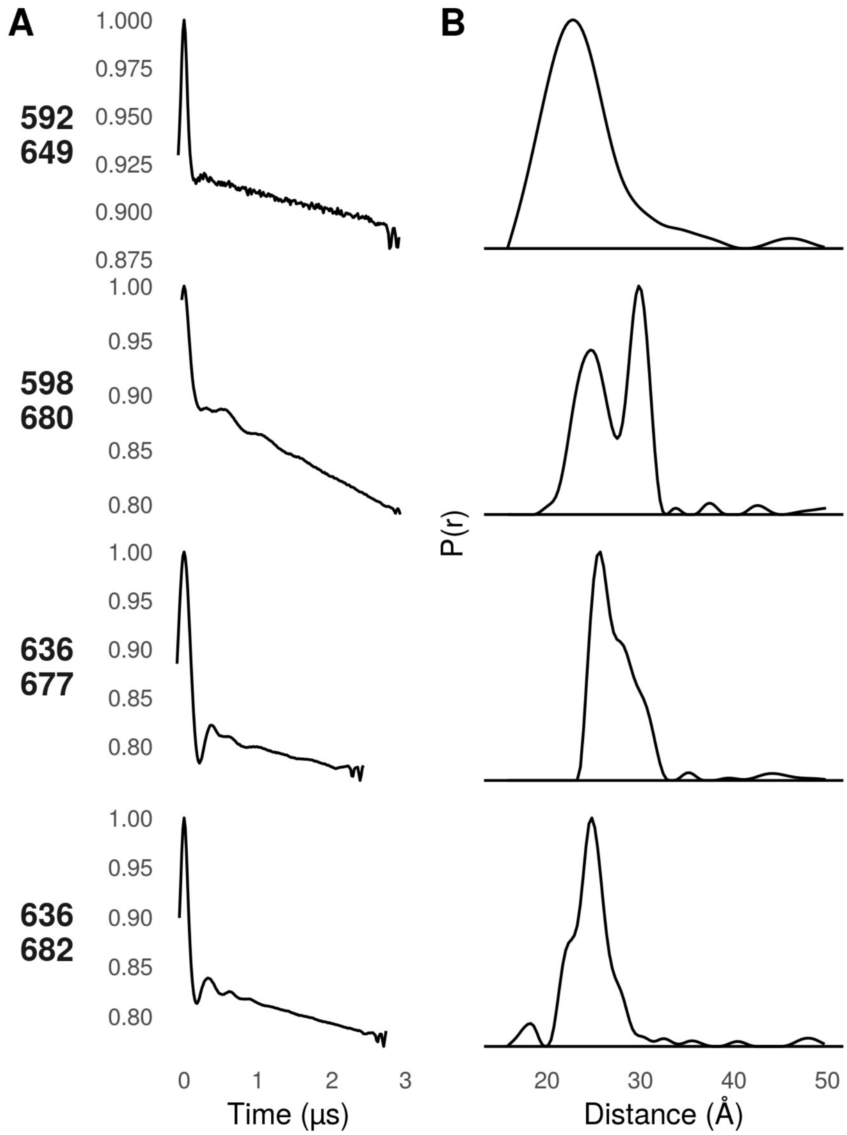


**Figure S1 . Data gathered in the ExoU C-terminus for this study.**

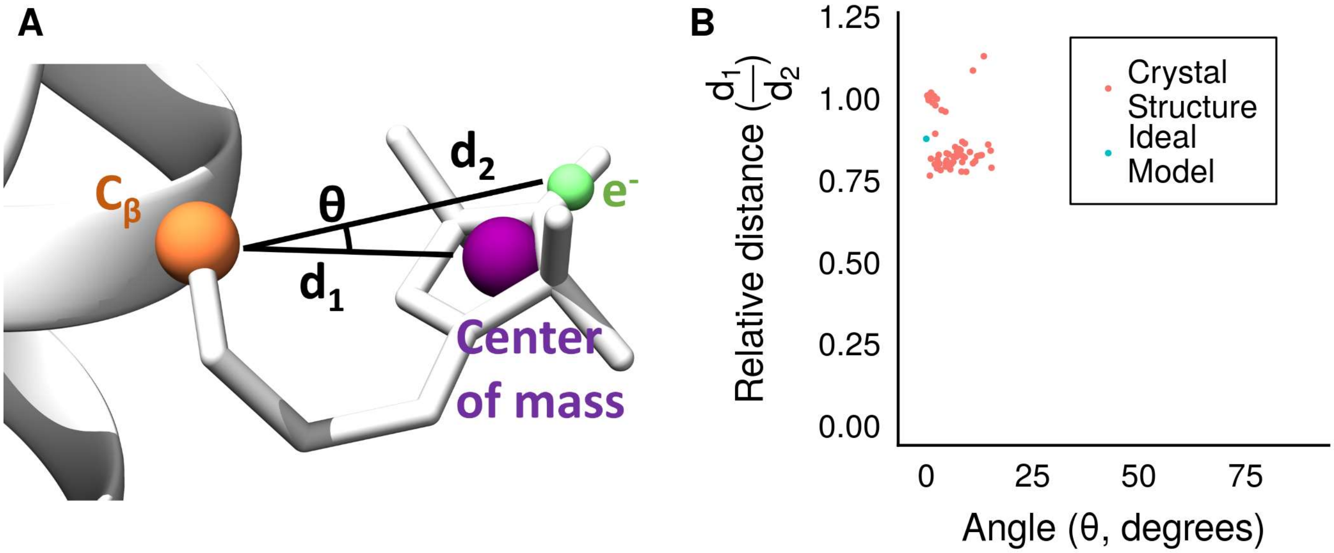


**Figure S2. Nitroxide centers of mass fall along the $C_\beta$-electron vector. A)** Depiction of spin label from PDB: 2Q9D showing the nitroxide center of mass (purple) and the nitroxide bond midpoint (green). **B).** Angle and relative distance of nitroxide center of mass along the $C_\beta$-electron vector.

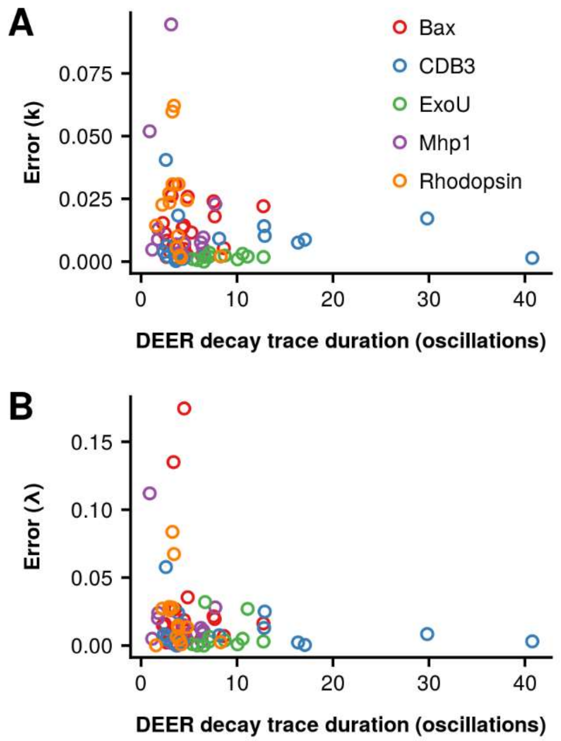


**Figure S3. Deviation between experimental and simulated background decay (*k*) and modulation depths (*λ*).** Oscillations were calculated from the average distance as previously described.

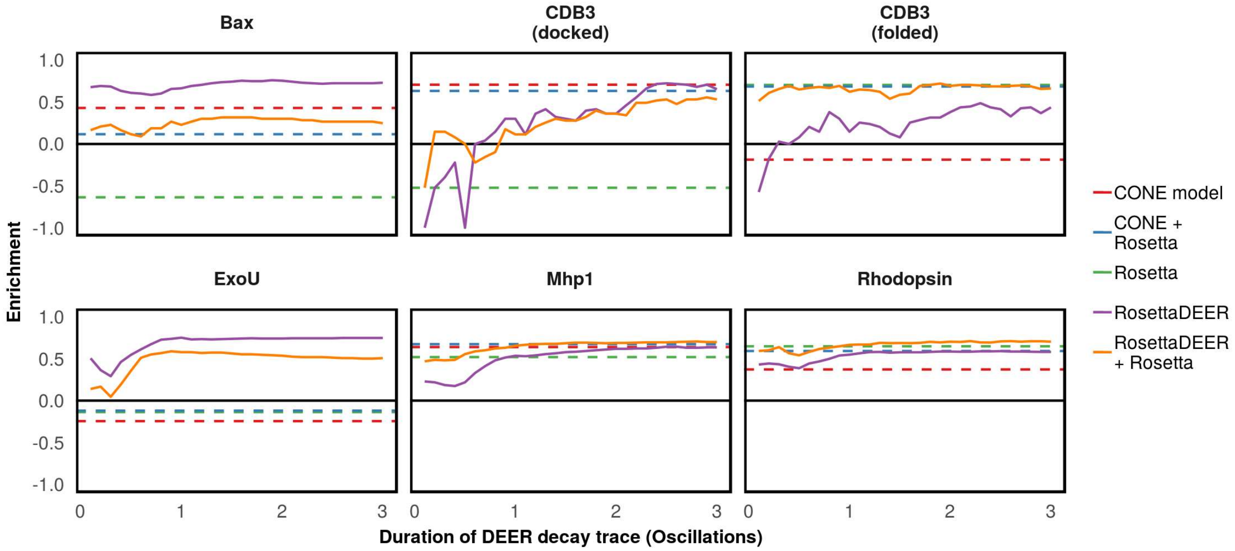


**Figure S4. Enrichment of misfolded and misdocked decoys as a function of DEER decay trace duration.** Enrichment was quantified as the logarithm of the percentage of native-like models (top 10% by RMSD100SSE) that were also in the top 10% by score. An enrichment of 1 indicates perfect correlation between model quality and score.

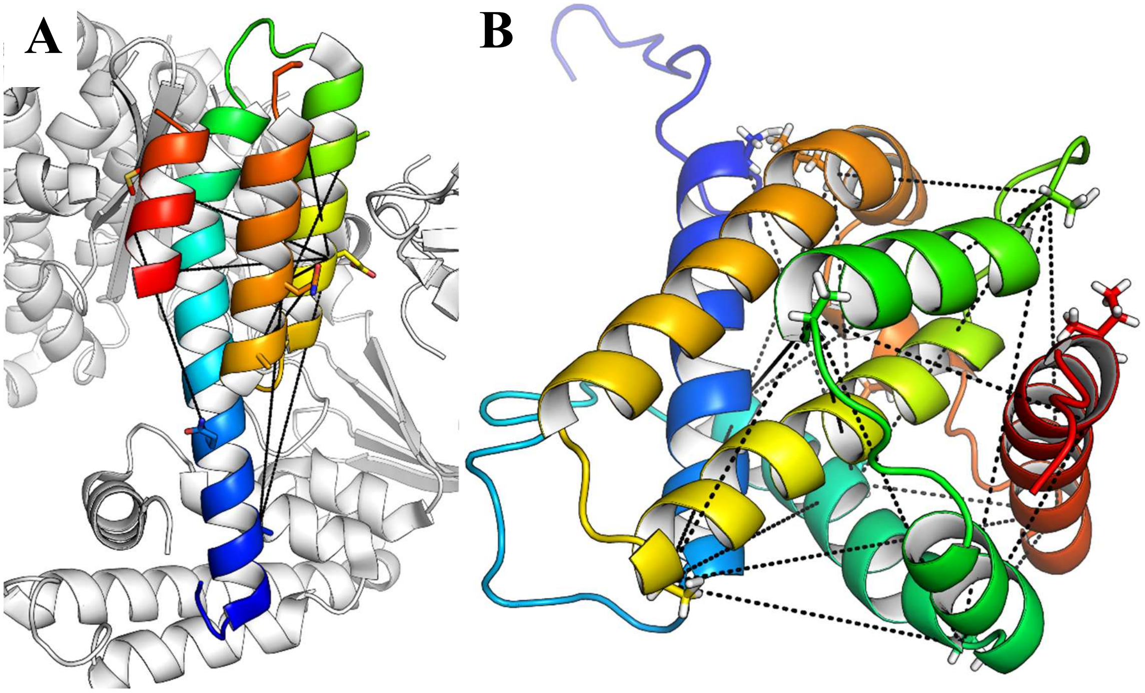


**Figure S5. Placement of experimental DEER restraints on ExoU (PDB: 3TU3, A) and Bax (PDB: 1F16, chain 8, B).** The N-terminus of ExoU, which was not modeled, is shown in white.

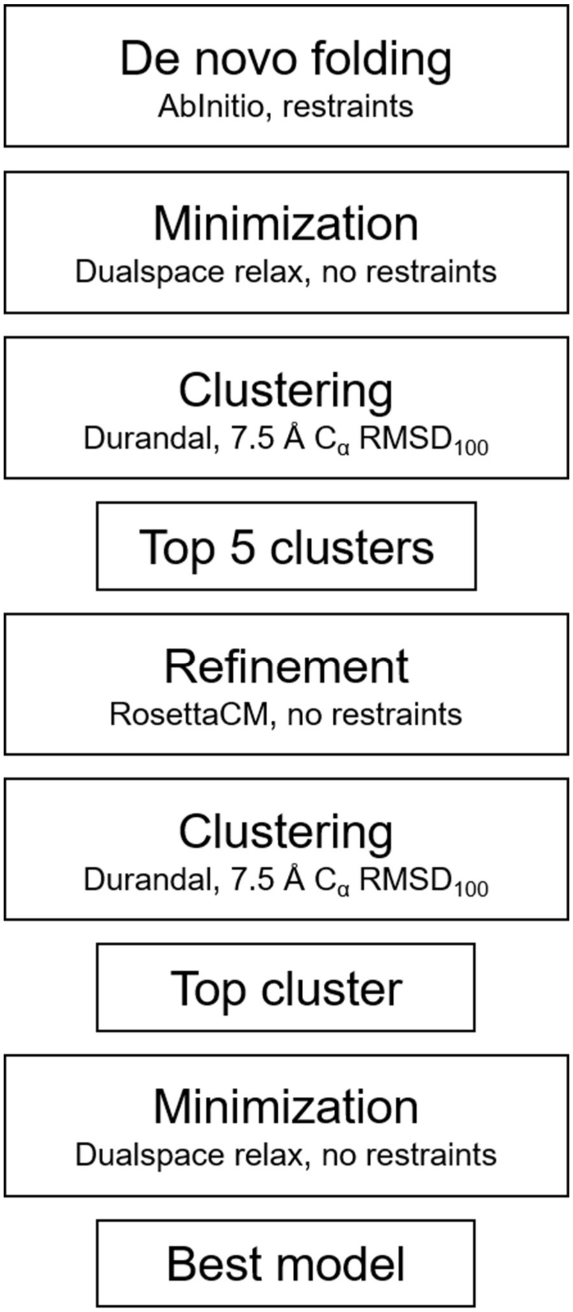


**Figure S6. Structure prediction pipeline for Bax and ExoU.**

40 Carugo O, Pongor S. A normalized root-mean-spuare distance for comparing protein

three-dimensional structures. *Protein Sci* 2002;**10**:1470–3. https://doi.org/10.1110/ps.690101.

41 Alford RF, Leaver-Fay A, Jeliazkov JR, O ’meara MJ, Dimaio FP, Park H, *et al.* The Rosetta all-atom energy function for macromolecular modeling and design 2017:1–35. https://doi.org/10.1101/106054.

42 Berenger F, Shrestha R, Zhou Y, Simoncini D, Zhang KYJ. Durandal: Fast exact clustering of protein decoys. *J Comput Chem* 2012;**33**:471–4. https://doi.org/10.1002/jcc.21988.

43 Shortle D, Simons KT, Baker D. Clustering of low-energy conformations near the native structures of small proteins. *Proc Natl Acad Sci* 1998;**95**:11158–62. https://doi.org/10.1073/pnas.95.19.11158.

44 Park H, Ovchinnikov S, Kim DE, DiMaio F, Baker D. Protein homology model refinement by large-scale energy optimization. *Proc Natl Acad Sci* 2018:201719115. https://doi.org/10.1073/pnas.1719115115.

45 Song Y, Dimaio F, Wang RYR, Kim D, Miles C, Brunette T, *et al.* High-resolution comparative modeling with RosettaCM. *Structure* 2013;**21**:1735–42. https://doi.org/10.1016/j.str.2013.08.005.

46 Conway P, Tyka MD, DiMaio F, Konerding DE, Baker D. Relaxation of backbone bond geometry improves protein energy landscape modeling. *Protein Sci* 2014;**23**:47–55. https://doi.org/10.1002/pro.2389.

47 Bleicken S, Jeschke G, Stegmueller C, Salvador-Gallego R, García-Sáez AJ, Bordignon E. Structural Model of Active Bax at the Membrane. *Mol Cell* 2014;**56**:496–505. https://doi.org/10.1016/j.molcel.2014.09.022.

48 Tessmer MH, Anderson DM, Buchaklian A, Frank DW, Feix JB. Cooperative substrate-cofactor interactions and membrane localization of the bacterial phospholipase A2(PLA2) enzyme, ExoU. *J Biol Chem* 2017;**292**:3411–9. https://doi.org/10.1074/jbc.M116.760074.

49 Jeschke G, Chechik V, Ionita P, Godt A, Zimmermann H, Banham J, *et al.* DeerAnalysis2006—a comprehensive software package for analyzing pulsed ELDOR data. *Appl Magn Reson* 2006;**30**:473–98. https://doi.org/10.1007/BF03166213.

50 Jo S, Cheng X, Islam SM, Huang L, Rui H, Zhu A, *et al.* CHARMM-GUI PDB manipulator for advanced modeling and simulations of proteins containing nonstandard residues. *Adv Protein Chem Struct Biol* 2014;**96**:235–65. https://doi.org/10.1016/bs.apcsb.2014.06.002.

51 Yarov-Yarovoy V, Schonbrun J, Baker D. Multipass membrane protein structure prediction using Rosetta. *Proteins* 2006;**62**:1010–25. https://doi.org/10.1002/prot.20817.

52 Viklund H, Elofsson A. OCTOPUS: Improving topology prediction by two-track ANN-based preference scores and an extended topological grammar. *Bioinformatics* 2008;**24**:1662–8. https://doi.org/10.1093/bioinformatics/btn221.

53 Hustedt EJ, Martinelli F, Stein RA, Faraldo-Gomez J, Mchaourab HS. Confidence Analysis of DEER Data and its Structural Interpretation with Ensemble-Biased Metadynamics. *Biophys J* 2018:1–17. https://doi.org/10.1101/299941.

54 Zhou Z, DeSensi SC, Stein RA, Brandon S, Dixit M, McArdle EJ, *et al.* Solution structure of the cytoplasmic domain of erythrocyte membrane band 3 determined by site-directed spin labeling. *Biochemistry* 2005;**44**:15115–28. https://doi.org/10.1021/bi050931t.

55 Altenbach C, Kusnetzow AK, Ernst OP, Hofmann KP, Hubbell WL. High-resolution distance mapping in rhodopsin reveals the pattern of helix movement due to activation. *Proc Natl Acad Sci* 2008;**105**:7439–44. https://doi.org/10.1073/pnas.0802515105.
